# Tuning gene expression to music: the compensatory effect of music on age-related cognitive disorders

**DOI:** 10.1101/2023.09.12.557408

**Authors:** Alberto Gómez-Carballa, Laura Navarro, Jacobo Pardo-Seco, Xabier Bello, Sara Pischedda, Sandra Viz-Lasheras, Alba Camino-Mera, María José Currás, Isabel Ferreirós, Narmeen Mallah, Sara Rey-Vázquez, Lorenzo Redondo, Ana Dacosta-Urbieta, Fernando Caamaño-Viña, Irene Rivero-Calle, Carmen Rodriguez-Tenreiro, Federico Martinón-Torres, Antonio Salas

## Abstract

Extensive literature has explored the beneficial effects of music in age-related cognitive disorders (ACD), but limited knowledge exists regarding its impact on gene expression. We analyzed transcriptomes of ACD patients and healthy controls, pre-post a music session (*n*=60), and main genes/pathways were compared to those dysregulated in mild cognitive impairment (MCI) and Alzheimer’s disease (AD) as revealed by a multi-cohort study (*n*=1269 MCI/AD and controls). Music was associated with 2.3 times more whole-genome gene expression, particularly on neurodegeneration-related genes, in ACD than controls. Co-expressed gene-modules and pathways analysis demonstrated that music impacted autophagy, vesicle and endosome organization, biological processes commonly dysregulated in MCI/AD. Notably, the data indicated a strong negative correlation between musically-modified genes/pathways in ACD and those dysregulated in MCI/AD. These findings highlight the compensatory effect of music on genes/biological processes affected in MCI/AD, providing insights into the molecular mechanisms underlying the benefits of music on these disorders.

## Introduction

What is the genuine impact of musical stimuli on human beings? Music, as an artistic manifestation, intricately blends sounds and silences to craft harmonious compositions encompassing elements of melody, timbre, pitch, and rhythm. It serves as a means of human expression and aesthetic communication, offering a profound impact on the emotional, cognitive, and physiological aspects of individuals’ experiences. In the realm of physics, sound is however, a vibration (mechanical disturbance) that propagates through the air or other media as an acoustic wave. The multifaceted influence of music on brain plasticity, autobiographical memory, and emotions represents a complex phenomenon that requires comprehensive scientific exploration. While numerous studies have been conducted in the field of neuroscience and cognitive sciences [1], there is still limited understanding of the underlying biochemical changes responsible for the observed effects. To bridge this gap, investigating genes and their expression under the stimulus of music constitutes an innovative approach to understanding the molecular mechanisms that underpin its beneficial effects on health. Despite the significant advancements in omics sciences over the past two decades, only four discrete attempts have been made to analyze the impact of music on transcriptomes [2]. Kanduri et al. [3] observed that exposure to 20 minutes of classical music led to the differential expression of 45 genes in musically experienced participants; however, no significant results were detected in musically inexperienced participants [4]; they also identified the involvement of *GATA2*, a gene previously linked to musical aptitude [5]. The same research group further examined the transcriptomic changes occurring in professional musicians by analyzing their pre- and post-performance transcriptomes following a 2-hour concert [6]; that study revealed up-regulated pathways related to dopaminergic neurotransmission, motor behavior, neuronal plasticity, and neurocognitive functions (e.g., learning and memory) [6]. Two further studies [7, 8] identified microRNAs that were up- and down-regulated of both listeners and professional musicians, building upon previous findings [3, 6]. Navarro et al. [1] used publicly available transcriptomic data from Alzheimer’s disease (AD) patients cross-compared to music-related genes and found a notable commonality of these genes with those altered in AD patients. None of all these studies analyzed the direct impact of music in age-related cognitive disease (ACD) patients.

The aforementioned studies shed light on the intricate relationship between music and gene expression [3-6]. However, further empirical research is needed to gain a comprehensive understanding of the specific molecular mechanisms underlying the association between music and gene expression [2]. The current investigation aligns with the concept of musical sensogenomics [2] and serves as the initial effort to analyze the impact of music on the transcriptomes of individuals with ACD. In contrast to previous attempts, this study has been meticulously designed to replicate an ecological environment, that is, resembling the natural or real-life conditions in which music is typically experienced by individuals, thereby minimizing the potential influence of confounding factors [2].

## Methods

### Participants, RNA isolation and RNA-Seq analysis

Under the frame of the Sensogenomics project (www.sensogenomics.com) and for the purpose of the present project, an experimental concert of classical music with a duration of 50 minutes was organized on June 14, 2022, in the ‘Auditorio de Galicia’ located in the city of Santiago of Compostela (Galicia, Spain). The audience included, among others, patients diagnosed with mild cognitive impairment (MCI) and dementia (henceforth referred to as the age-related cognitive disorder [ACD] cohort), and a healthy control cohort; **Supplementary Text**.

We collected blood samples using a procedure of capillary puncture before (Time Point 1 or TP1) and after (Time Point 2 or TP2) exposure to music (**Figure 1**), including 16 ACD patients (aged 64–93, mean 81.8), and 14 healthy donors (aged 18–74; mean 48.2]; **Table 1**. Sample collection was adapted to administratively accessible population and procedures. RNA was isolated using the PAXgene blood miRNA extraction kit (Qiagen) adapting buffer volumes to the amount of blood collected (100–200μl). Sequencing was performed using a NovaSeq 6000 System (100 paired end; ∼70M reads/ sample); **Supplementary Text**.

**Figure 1.**
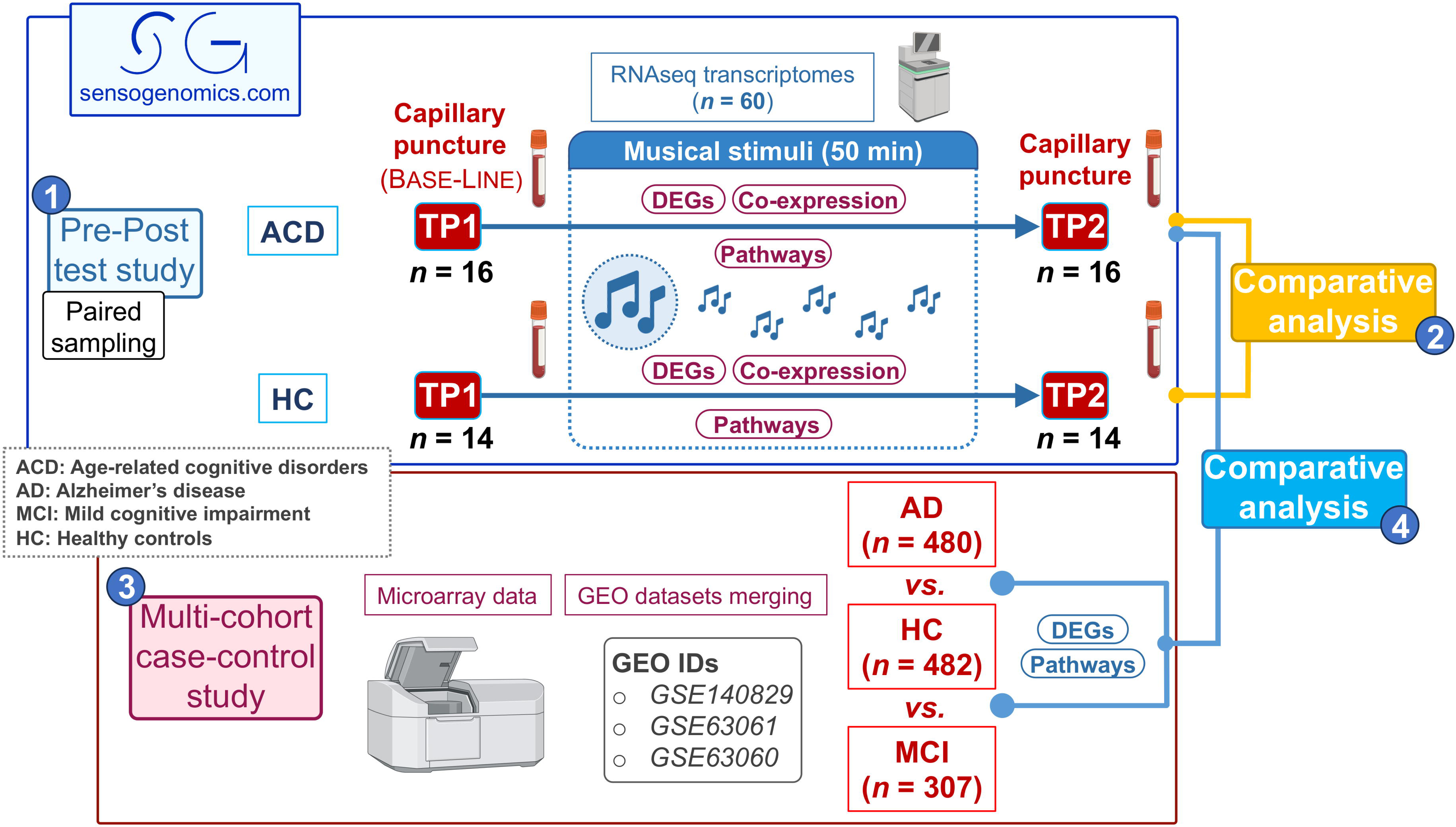
Scheme showing the overall experimental and analytical design. Numbers in circles indicate the four different blocks of analysis carried out in the present study: 1) Pretest-Posttest test study comparing transcriptomes before and after listening to music (same subjects) from ACD patients and healthy controls separately; 2) comparative analysis between transcriptomic response to music in ACD and healthy donors; 3) multi-cohort case-control study to detect MCI/AD-related genes and pathways; and 4) comparative analysis to study the impact of the musical stimuli on the expression of genes and pathways altered in MCI/AD patients.

**Table 1.** Demographic characteristics of donors.

|  | Healthy controls | ACD patients |
| --- | --- | --- |
| **Gender** |  |  |
| *Female* | 12 (85.7%) | 10 (62.5%) |
| *Male* | 2 (14.3%) | 6 (37.5%) |
| **Age** |  |  |
| *Mean (SD)* | 48.2 (18.0) | 81.8 (9.2) |
| *Range* | 18-74 | 64-93 |
| **Disease condition** |  |  |
| *MCI* | – | 8 (50.0%) |
| *AD* | – | 2 (12.5%) |
| *Non-AD dementia* | – | 6 (37.5%) |

### Statistical analysis

*DESeq2* [9] R package was used to normalize counts and carry out the differential expression (DE) analysis after removing lowly expressed genes. We first carried out a pre-post study design by comparing samples collected before the musical stimuli (baseline; TP1) to samples collected after the musical stimuli (TP2) from the same healthy controls and ACD patients. To isolate transcriptomic changes exclusively related to the musical stimuli, we used a paired-samples design accounting for patient-to-patient differences with respect to TP1. For the Principal Component Analysis (PCA), we defined the transcriptome displacement (*TD*) score to quantitatively measure the ‘global displacement of transcriptomes’ from TP1 to TP2 as reflected in the Euclidian space of the plot. *TD* is the mean of the absolute values of the differences between individual *i* projections from TP1 and TP2 coordinates on the *j* different principal components (e.g., PC1 and PC2); *TD* is computed for the two cohorts, e.g., in ACD patients: *TD_ACD_*_,*i*,*j*_=mean (|TP1*_ACD,i,j_* – TP2*_ACD,i,j_*|). We additionally estimated the specific weight of the main genes driving the variability in PC1 (top and bottom 10% of the loading range in PC1) using *PCAtools* [10].

In a different comparative analysis, we evaluated the effect of listening to music in ACD patients with differentially expressed genes (DEGs) and pathways (DEPs) that are altered in MCI and AD patients when these are compared to controls. For this purpose, data from three independent microarray datasets analyzing MCI and AD patients and healthy controls were downloaded from Gene Expression Omnibus (GEO) accession numbers GSE140829, GSE63061 and GSE63060 (case-control study). By using this multi-cohort strategy, we aimed at having a statistically robust age-matched group of AD/MCI patients and controls to contrast them with our findings. After an initial quality control of each dataset separately, and the removal of outliers, the whole multi-cohort dataset was composed of gene expression data from 307 MCI patients (mean age=75.1 [SD=7.2]), 480 AD patients (mean age=75.1 [SD=7.1]) and 482 age-matched healthy controls (mean age=73.8 [SD=6.3]) (**Figure 1**). Processing and merging of the data as well as the case-control DE analysis between AD and controls was carried out as explained elsewhere [1]. STRING database [11] was used to build the Protein-Protein Interaction (PPI) network of the shared DEGs obtained from AD *vs.* controls (multi-cohort study), and from TP2 *vs.* TP1 in ACD patients (Pretest/Posttest study). Visualization and pathways functional enrichment of the PPI was carried out with Cytoscape software [12].

Biological pathways were inferred directly from gene expression data using Gene Set Variation Analysis (*GSVA*) R package [13] and Quantitative Set Analysis for Gene Expression (*QuSAGE*) R package [14]; see details in **Supplementary Text**. In *QuSAGE*, to explore the most differentially activated pathways between ACD and healthy donors in response to music, we consider conservative filters of pathway activity (|PA| in ACD + |PA| in controls > 0.05) and statistical significance (*P*-value < 0.05). Heatmaps and rain cloud plots were built using *EnhancedVolcano* [15], *ComplexHeatmap* [16] and *Raincloudplots* [17], respectively. The Wilcoxon test was used to assess statistical significance between patient groups, and the Spearman test (ρ representing the Spearman’s coefficient) was used to compute correlation values. We investigated clusters of co-expressed genes potentially correlated to the musical stimuli in both the ACD and control groups, separately. Gene expression data normalized and corrected for patient-to-patient differences was used to build a signed weighted correlation network with the Weighted Gene Co-expression Network Analysis (*WGCNA*) R package [18]. Only genes that showed the most variant expression values between samples (the top 75% with the highest variance) were included in the analysis. See **Supplementary Text** for more information on co-expression analysis.

Adjustment for multiple test was carried out using the FDR method by Benjamini-Hochberg [19] (henceforth ‘FDR’ refers to adjusted *P*-value). All statistical analyses were performed using the statistical software R v. 4.2.2. [20].

## Results

### Global impact of music on the human transcriptome

We first aimed at quantifying the global effect of music on the transcriptomes of the two groups of donors separately. ACD patients exposed to music showed 2.3 times more DEGs (*n*=2,605) than controls (*n*=1,148); **Table 2**. Moreover, while the proportion up-regulated/down-regulated DEGs was balanced in ACD patients (1.315/1.290=1.02), it was significantly lower in controls (528/620=0.85). This differential pattern of gene expression is very clear in a density plot of fold-change (FC) values of the DEGs in both groups (**Figure 2A**): the median in ACD patients is displaced towards positive FC values (0.10) compared to the median in controls (-0.11). Among the DEGs, the proportion of protein coding genes was higher in ACD patients than in controls (1.89 in ACD patients *vs*. 1.24 in controls); **Table 2**.

**Figure 2.**
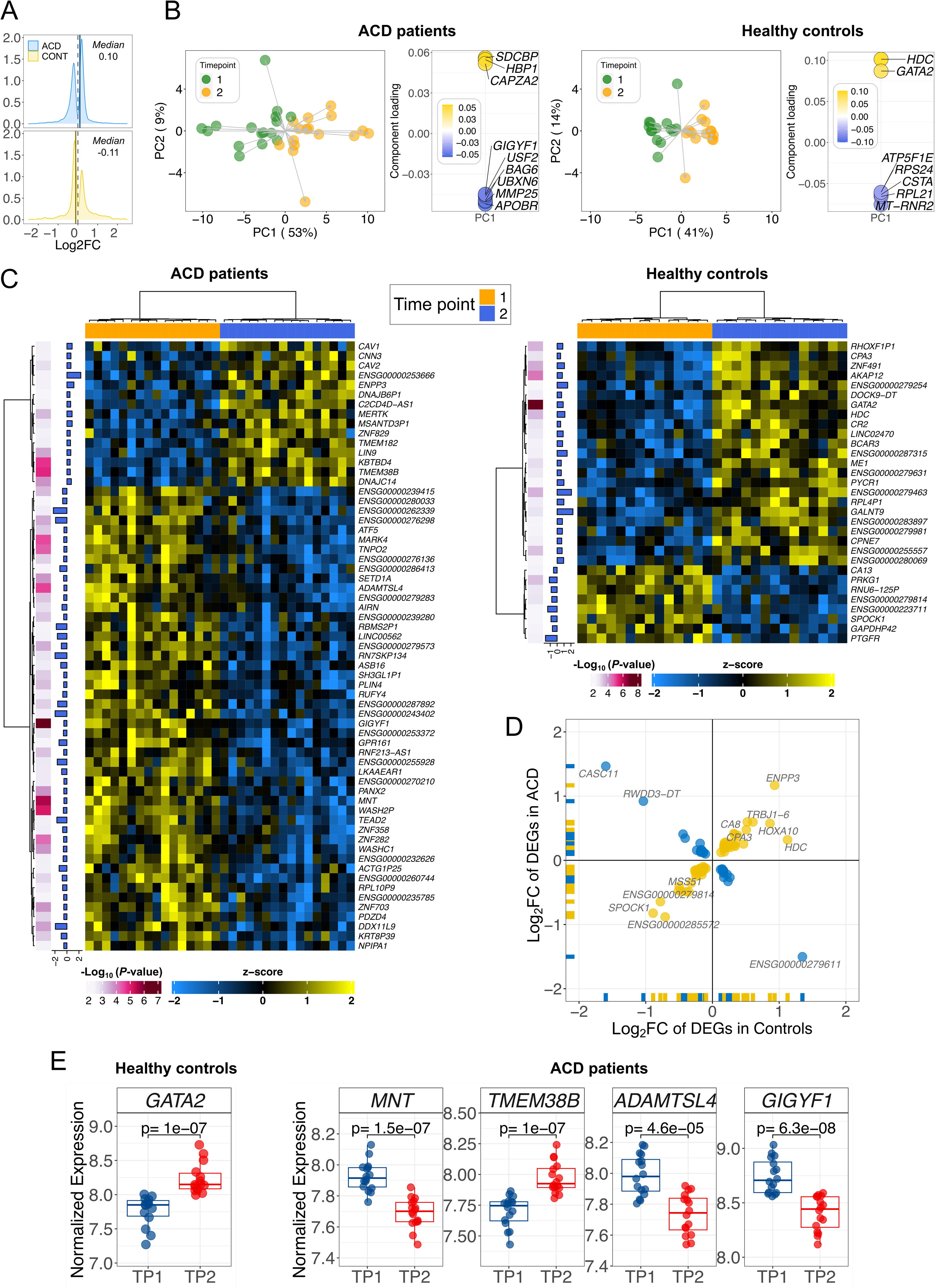
Global transcriptome patterns and differential expression analysis in ACD and healthy controls. (A) Density plots of log_2_FC; (B) Principal Component Analysis (left) and the genes that play the main role in the topology of the PC1 (right). (C) Heatmaps of the most remarkable DEGs (*P*-values < 0.005 and |log_2_FC|>0.5) in ACD patients and healthy controls when contrasting the two time points (A). (D) Correlation of DEGs in the two cohorts. (C) DEGs showing the most extreme and significant differences in ACD patients and controls.

**Table 2.** Biotypes of the DEGs observed in the control and the ACD patient cohorts. UR: up-regulated; DR: down-regulated.

| Biotype | Controls |  |  | ACD patients |  |  |
| --- | --- | --- | --- | --- | --- | --- |
|  | *n* | *n*<sub>UR</sub> | *n*<sub>DR</sub> | *n* | *n*<sub>UR</sub> | *n*<sub>DR</sub> |
| **IG C gene** | 2 | 2 | – | – | – | – |
| **IG V gene** | 1 | – | 1 | 5 | 1 | 4 |
| **IG V pseudogene** | 1 | 1 | – | – | – | – |
| **lncRNA** | 140 | 81 | 59 | 243 | 60 | 183 |
| **miRNA** | 1 | 1 | – | – | – | – |
| **Misc RNA** | 3 | 2 | 1 | 9 | – | 9 |
| **Mt rRNA** | 1 | – | 1 | – | – | – |
| **Processed pseudogene** | 37 | 21 | 16 | 52 | 12 | 40 |
| **Protein coding** | 921 | 392 | 529 | 2,192 | 1,204 | 988 |
| **rRNA pseudogene** | – | – | – | 2 | – | 2 |
| **snoRNA** | 1 | 1 | – | – | – | – |
| **snRNA** | 4 | 1 | 3 | 3 | 1 | 2 |
| **TEC** | 14 | 9 | 5 | 32 | 1 | 31 |
| **TR C gene** | – | – | – | 2 | 2 | – |
| **TR J gene** | 1 | 1 | – | 2 | 2 | – |
| **TR V gene** | – | – | – | 12 | 12 | – |
| **Transcribed processed pseudogene** | 2 | 1 | 1 | 11 | 3 | 8 |
| **Transcribed unitary pseudogene** | 3 | 3 | – | 4 | 2 | 2 |
| **Transcribed unprocessed pseudogene** | 11 | 9 | 2 | 20 | 12 | 8 |
| **Translated processed pseudogene** | – | – | – | 1 | 1 | – |
| **Unitary pseudogene** | – | – | – | 2 | – | 2 |
| **Unprocessed pseudogene** | 4 | 3 | 1 | 9 | 1 | 8 |
| **No data** | 1 | – | 1 | 4 | 1 | 3 |
| **Total** | 1,148 | 528 | 620 | 2,605 | 1,315 | 1,290 |

PCA (**Figure 2B**) showed the displacement of transcriptomes in TP2 with respect to TP1, revealing also the remarkable differences existing between ACD donors and healthy controls. The main pattern emerging from these plots is the clear differentiation existing between time points; especially marked in the first principal component (PC1; accounting for 53% and 41% of the variation in ACD patients and controls, respectively). The genes *GATA2* (in controls) and *GIGYF1* (in ACD patients) showed the strongest contribution to PC1, suggesting a key role in the observed differentiation between the groups at TP2. (**Figure 2B**). The displacement of individual transcriptomes was more pronounced in ACD patients than in controls (PC1: *TD_ACD_*_;1_=7.5; *TD_CO_*_;1_=4.6; PC2: *TD_ACD_*_;2_=2.5 [9% of the variation]; *TD_CO_*_;2_=1.8 [14%]). In good agreement with the PCA, the heatmap representations of DEGs (*P*-values < 0.005 and |log_2_FC| > 0.5) show the main differentiation in both cohorts comes from TP1 *vs*. TP2 (**Figure 2C).** Heatmap representations of DEGs confirmed that the main differentiation between ACD patients and controls occurred between TP1 and TP2 (**Figure S1)**. A comparison of the common genes that are DEGs in the two cohorts, indicates that most of them (59/84; 70%) show a positively correlated expression pattern (ρ=0.95, *P*-value=2.2×10^-06^), and only a few genes (25/84; 30%) appear to be negatively correlated (ρ=0.98, *P*-value=2.2×10^-06^); therefore, resulting in a positive overall correlation value (ρ=0.5, *P*-value=2.2×10^-06^); **Figure 2D**.

While there is only one gene surpassing the multiple test threshold of significance in healthy controls, namely, *GATA2* (up-regulated in TP2; *P*-value=1.0×10^-7^), there are 328 genes surpassing the multiple test threshold in ACD cases. Among the DEGs that show the most extreme and significant expression changes before and after musical stimuli in ACD patients are the down-regulated genes *GIGYF1* (*P*-value=6.3×10^-8^), and *MNT* (*P*-value=1.5×10^-7^), *ADAMTSL4* (*P*-value=4.6×10^-5^) and the up-regulated gene *TMEM38B* (*P*-value=1.0×10^-7^) (**Figure 2E**). A full list of significant DEGs is provided in **Table S1**.

### Music effect on key neurological-related pathways

GSVA was carried out to better understand the main biological pathways representing the differences observed between the expression of genes in patients and controls when exposed to music. GSVA yielded 556 DEPs in ACD patients, and 512 DEPs in healthy controls, 49 and 38 of them were related to neurobiological processes in ACD patients and controls, respectively (**Table S2**). Interestingly, we found that music impacts key pathways so strongly that the patterns generated allow to perfectly discriminate transcriptome profiles by their time points (see heatmaps in **Figure S2**). Most of the statistically significant DEPs involved in neuro-biological processes are exclusive to one of the two cohorts analyzed (see details in **Table S2**; **Figure S2**).

Remarkably, the top DEPs observed in the music-stimulated Alzheimer’s disease (ACD) cohort are closely related to pathways that are typically dysregulated in these neurodegenerative disorders. This observation is particularly noteworthy if we bear in mind that the DEPs were inferred using a paired-sampling design, which essentially measures the displacement of the transcriptome from TP1 to TP2. Among these pathways are the over-activated “regulation of amyloid beta clearance” (log_2_FC=0.24; FDR=0.0095), “L glutamate import across plasma membrane” (log_2_FC=0.43; FDR=6.1×10^-7^), and the “sphingomyelin biosynthetic process” (log_2_FC=0.33; FDR=0.0099); but also the under-activated “positive regulation of glutamate secretion” (log_2_FC=-0.34; FDR=0.004) (**Figure S2**). In addition, we have observed that the pathway “dopamine biosynthetic process” was also activated by the musical stimuli in ACD donors (log_2_FC=0.41; FDR=0.0014).

In controls, there are a few DEPs that are closely related to neurological processes and neurotransmission, such as the down-regulated “negative regulation of oxidative stress induced neuron death” (log_2_FC=-0.33; FDR=0.0001), “neuron death in response to oxidative stress” (log_2_FC=-0.24; FDR=0.0033), and the up-regulated “regulation of neurotransmitter transport” (log_2_FC=-0.16; FDR=0.0081) (**Figure S2**). Nevertheless, it is also noticeable that the activation of the pathway “L glutamate import across plasma membrane”, observed in ACD patients in response to music, stands in contrast with the under-activation observed in healthy controls (log_2_FC=-0.25; FDR=0.0045); **Figure S3; Table S2**.

QuSAGE allows to evaluate the pathways showing the most extreme differences in activation (pathway activation; PA) between patients and controls following the musical stimuli; **Table S2** and **Figure S4A**. As in the case of GSVA analysis, the observed measurements from this paired-sampling QuSAGE analysis (TP1 *vs*. TP2), must be attributed primarily to the action of the musical stimuli impacting the phenotype condition (ACD and healthy controls).

In good agreement with GSVA, the pathway that shows the highest *PA* in ACD patients is “negative regulation of amyloid beta clearance”, while it is virtually inactive in healthy controls (*PA_ACD_*_=_0.14; PA_HC_=0.01; *P*-value=0.0032; **Figure S4B**). Another pathway showing major differentiation between these groups is “regulation of L glutamate import across plasma membrane”, being the second most activated in ACD patients (*PA_ACD_*=0.14; *PA_HC_*=-0.06; *P*-value=0.0025); its related pathway “L-glutamate import across plasma membrane” shows even a more statistically significant difference between ACD patients and controls (*PA_ACD_*=0.09; *PA_HC_*=-0.05; *P*-value=0.0016), **Table S2**. QuSAGE also reveals important differences in the impact of the musical stimuli between ACD patients and controls in dopamine related pathways: the “dopamine biosynthetic process” is significantly more activated in ACD than in controls (*PA_ACD_*=0.083; *PA_HC_*=-0.01; *P*-value=0.027), the same applies to the “positive regulation of dopamine receptor signaling pathway” (*PA_ACD_*=0.12; *PA_HC_*=-0.03; *P*-value=0.039); **Table S2**. Also interesting is the recurrent presence of pathways related to sphingolipids metabolism: “cellular sphingolipid homeostasis” (*PA_ACD_*=-0.06; *PA_HC_*=0.04; *P*-value=0.005), “regulation of sphingolipid biosynthetic process” (*PA_ACD_*=-0.02; *PA_HC_*=0.07; *P*-value=0.005), “positive regulation of sphingolipid biosynthetic process” (*PA_ACD_*=-0.03; *PA_HC_*=0.13; *P*-value=0.0076), “sphingomyelin biosynthetic process” (*PA_ACD_*=0.08; *PA_HC_*=-0.02; *P*-value=0.0098), “sphingomyelin translocation” (*PA_ACD_*=-0.11; *PA_HC_*=0.04; *P*-value=0.013), and sphingomyelin catabolic process (*PA_ACD_*=-0.04; *PA_HC_*=0.07; *P*-value=0.042); **Table S2**.

Other notable neurology-related pathways showing important differences in expression between patients and controls are: “negative regulation of oxidative stress induced neuron death” (*PA_ACD_*=0.05; PA_HC_=-0.05; being the most statistically significant value, namely, *P*-value=0.0001), “trans synaptic signaling modulating synaptic transmission” (*PA_ACD_*=0.07; *PA_HC_*=-0.04; *P*-value=0.0063), “neuronal action potential propagation” (*PA_ACD_*=-0.009; *PA_HC_*=0.04; *P*-value=0095), and “neuron death in response to oxidative stress” (*PA_ACD_*=0.03; *PA_HC_*=-0.03; *P*-value=0.002).

### Co-expression modules driven by music modulate autophagy in ACDs

After filtering out genes with lower variability, a co-expression network analysis was conducted using the ACD and the control datasets (4,283 and 4,250 genes, respectively).

Interestingly, the WGCNA analysis revealed 22 clusters of co-expressed genes from ACD gene expression data, with 9 of them significantly correlated with the musical stimuli (**Figure 3A; Figure S5A; Figure S5B**). Multiple test correction was applied (FDR < 0.05), resulting in 7 statistically significant modules (**Table S3**), 3 of them positively correlated (*RRM1* [royalblue], *TPM3* [saddlebrown] and *CMTM6* [magenta]) and 4 negatively correlated (*TRIM56* [black], *UPF1* [darkmagenta], *PRKD2* [paleturquoise] and *ARFGAP1* [cyan]) with the music stimulus. This global over-activation or under-activation in TP2 compared to TP1 is also notable when representing eigengene values for individual samples; the results are consistent with the correlation values, with modules *PRKD2*, *RRM1*, *TPM3*, *TRIM56* and *UPF1* exhibiting the most significant differences in individual expression patterns (**Figure 3B**). The *TRIM56* module showed the highest correlation (negative) with the musical stimuli (R=-0.70) and the lowest *P*-value (7.3×10^-6^), followed by *TPM3* (R=0.63; *P*-value=1.0×10^-4^) and *RRM1* (R=0.63; *P*-value=1.0×10^-4^); both indicating a global activation pattern. The influence of core genes, measured as module membership (MM), was evaluated, and 6 out of the 7 modules showed a high statistically significant correlation between individual gene MM and individual gene correlation with musical stimuli, indicating functional relatedness; the strongest link was detected for the *TPM3* (R=0.56; *P*-value=1.5×10^-63^) and *TRIM56* (R=0.83; *P*-value=1.2×10^-55^) modules (**Figure S5C**).

**Figure 3.**
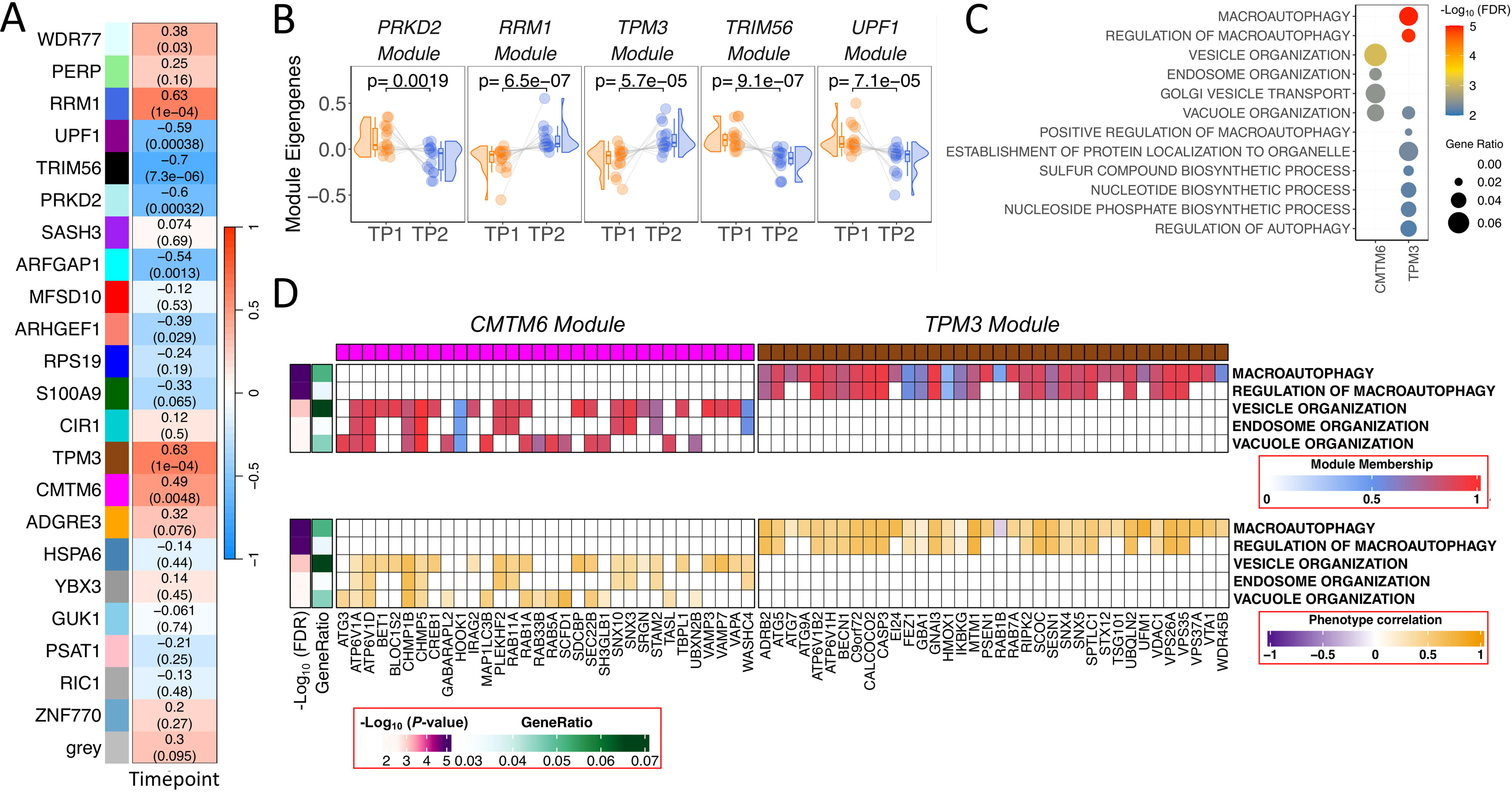
Co-expression analysis in ACD patients. (A) Correlation values heatmap obtained from WGCNA module analysis of co-expressed genes in ACD patients. Upper value corresponds to the individual correlation value of the module with musical stimuli. *P*-values of these correlations are shown in brackets (B) Raincloud plots of differences in individual samples eigengenes between TP1 and TP2 from modules showing the largest statistical significance in ACD patients. (C) Top biological processes detected from most statistically significant modules. Modules with significant pathways are named as the hub gene names *CMTM6* and *TPM3*. (D) Module membership (MM) and phenotype correlation (musical stimuli) for genes from the most significant pathways obtained from the modules *CMTM6* and *TPM3*.

Pathway analysis of the genes in the modules identified statistically significant pathways in modules ‘*TPM3’* and ‘CMTM6’ (**Table S3; Figure 3C**). The ‘*TPM3’* module was associated with macroautophagy (FDR=1.15×10^-5^), while ‘*CMTM6’* was linked to vesicle, endosome and vacuole organization (FDR=0.003). Shared genes between these modules and pathways have higher MM values and showed a high correlation with the musical stimuli, indicating functional relatedness, as also suggested by the proximity observed for the pathways in the hierarchical clustering tree (**Figure 3C, 3D, S5B**).

In the control dataset, modules of co-expressed genes were not found to be statistically significant after multiple test correction (see **Supplementary Text** for additional results).

### Music elicits expression in the opposite direction to MCI/AD in patients

We cross-compared the genes and pathways emerging from the contrast ‘MCI *vs*. controls’, with the genes and pathways emerging from the comparison ‘TP1 *vs*. TP2’ in ACD patients; this comparison yielded 16,743 overlapping genes (FDR significant DEGs = 207; henceforth DEGs_MCI_) and 5,350 overlapping pathways (FDR significant DEPs = 227; henceforth DEPs_MCI_) (**Figure 4A**). We also conducted a cross-comparison of DEGs and DEPs that are altered in AD patients (compared to controls) with those emerging from musical stimuli in ACD patients. The analysis revealed 16,721 overlapping genes, including 131 FDR significant DEGs (DEGs_AD_), and 5,352 overlapping pathways, including 220 FDR significant DEPs (DEGs_AD_); **Figure 5A**.

**Figure 4.**
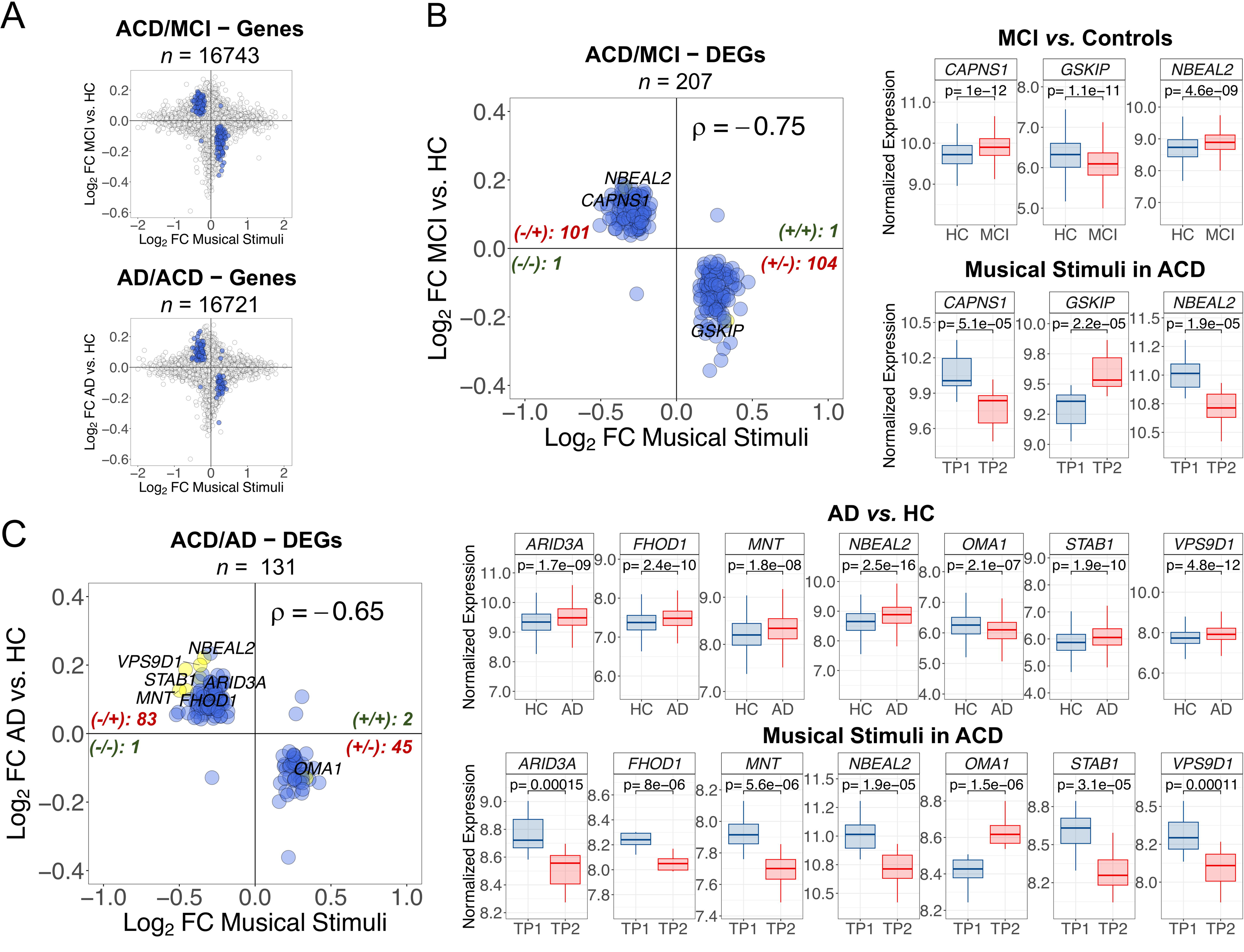
Correlation patterns of genes between the comparisons ‘ACD patients in TP1 *vs*. TP2’ to *i*) ‘MCI *vs*. healthy controls’, and *ii*) ‘AD *vs*. healthy controls’. (A) Dots indicate log_2_FC values for genes shared between the comparisons; blue dots are DEGs with significant FDR < 0.05. (B) FDR DEGs_MCI_; (C) FDR DEGs_AD_. For (B) and (C): only top DEG names are displayed (yellow points); these top elements were identified by first selecting the best 50 FDR values and top 50 higher |log_2_FC| from each comparison separately; next selecting the shared elements between the two comparisons. For the top DEGs we also show boxplots of (normalized) expression. Spearman correlation values and the number of genes plotted in each quadrant are indicated. For all the correlation values, *P*-value < 2.22×10^-16^. MCI: Mild Cognitive Impairment; AD: Alzheimer’s disease; HC: healthy controls; ACD: age-related cognitive disorder.

**Figure 5.**
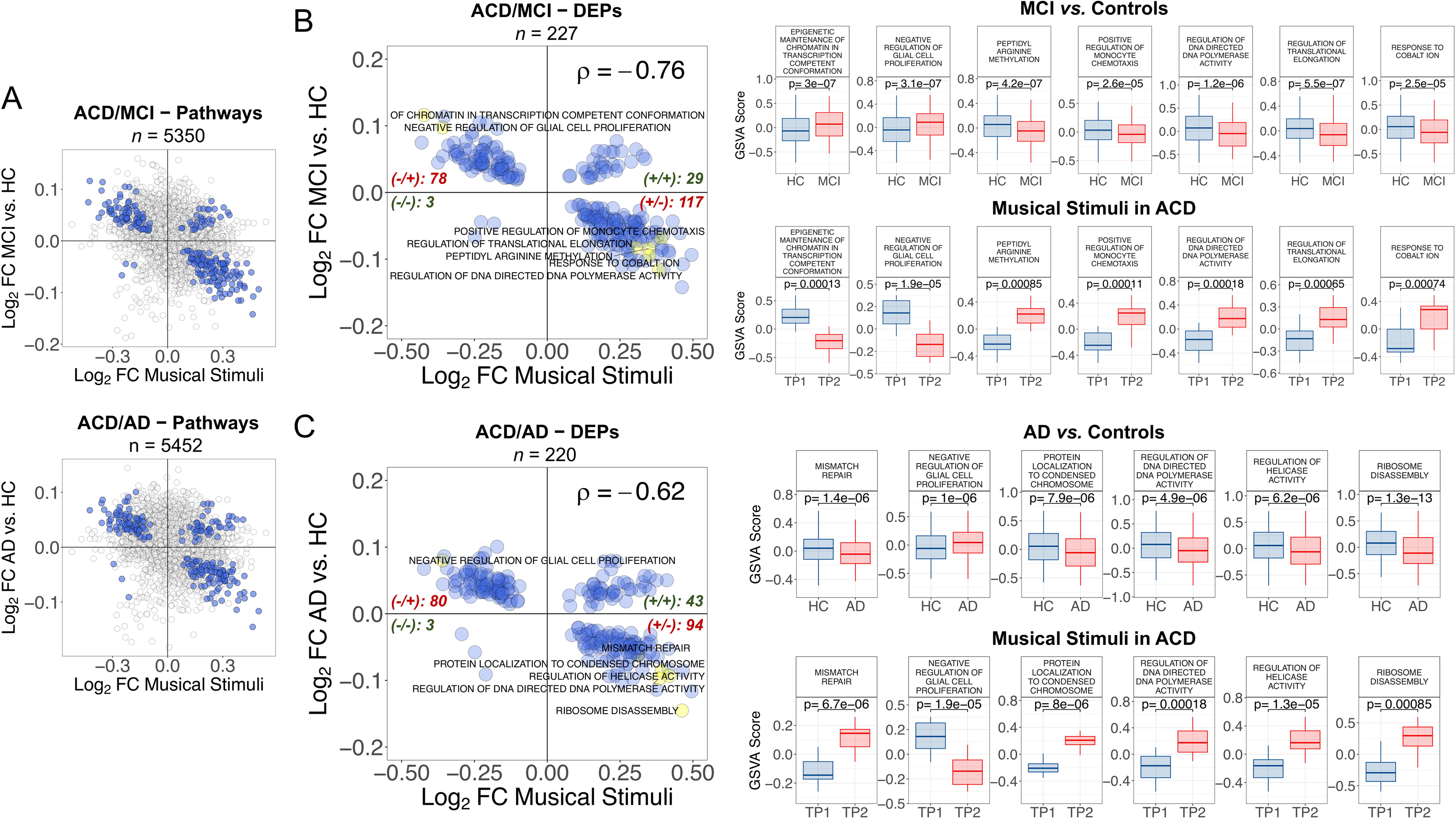
Correlation patterns of pathways between the comparisons ‘ACD patients in TP1 *vs*. TP2’ to *i*) ‘MCI *vs*. healthy controls’, and *ii*) ‘AD *vs*. healthy controls’. (A) Dots indicate log_2_FC values for pathways shared between the comparisons; blue dots are DEPs with significant FDR < 0.05. (B) FDR DEPs_MCI_; (C) FDR DEPs_AD_. For (B) and (C): only top DEP names are displayed (yellow points); these top elements were identified by first selecting the best 50 FDR values and top 50 higher |log_2_FC| from each comparison separately; next selecting the shared elements between the two comparisons. For the top DEGs we also show boxplots of *GSVA* score. Spearman correlation values and the number of genes plotted in each quadrant are indicated. For all the correlation values, *P*-value < 2.22×10^-16^. MCI: Mild Cognitive Impairment; AD: Alzheimer’s disease; HC: healthy controls; ACD: age-related cognitive disorder.

Of the total FDR DEGs_MCI_, 99% of them (205/207) showed log_2_FC that are negatively correlated with musically-stimulated DEGs (*n*=205; ρ=-0.65, *P*-value=2.2 × 10^-16^), the overall correlation being significantly negative (*n*=207; ρ=-0.75, *P*-value=2.2 × 10^-16^); **Figure 4B, Table S4**. Two of the most remarkable genes were up-regulated in MCI (*CAPNS1* and *NBEAL2*), and one was down-regulated (*GSKIP*); **Table S4**. The PPI-network and downstream enrichment analysis of the proteins coded by the top 207 genes (**Figure S6A; Table S4**) show interesting biological connections between these genes; these connections are more clearly visible in the enrichment map (**Figure S6A**). Thus, the most interesting biological process emerging from these genes is the “regulation of macroautophagy” (FDR=0.0057); **Figure S6A**, **Table S4**.

As seen with the MCI comparison, we observed a major proportion of FDR DEGs_AD_ (97.7%; 128/131) with negatively correlated log_2_FC (*n*=128; ρ=-0.68, *P*-value=2.2 × 10^-16^); the overall correlation being also negative (*n*=131; ρ=-0.65, *P*-value=2.2 × 10^-16^); **Figure 4C**. The most remarkable DEGs_AD_ were all up-regulated in AD (*FHOD1*, *MNT*, *STAB1*, *NBEAL2*, *ARID3A* and *VPS9D1*), except for one (*OMA1*), which showed down-regulation in AD with respect to control samples (*OMA1*); **Table S4**. We further explored the PPI-network and downstream enrichment analysis of the proteins coded by these top 131 genes (**Figure S6B; Table S4**); the enrichment map of this network (**Figure S6B**) indicates that the more important processes related to these genes are inter-connected within pathways related to G-alpha receptors (FDR=0.0361), AD (FDR=0.0063), and NRAGE cell death (FDR=0.0361 and 0.0417); **Table S4**.

The compensatory effect of the musical stimuli on the expression of genes involved in MCI is also evident when comparing log_2_FC of DEPs_MCI_ compared to musically-stimulated DEPs (**Figure 5B, Table S4**). A main proportion of the FDR shared DEPs showed negatively correlated log_2_FC values (85.9%; *n*=195; ρ=-0.90, *P*-value=2.2 × 10^-16^); the global log_2_FC correlation pattern being therefore negative (*n*=227; ρ=-0.76, *P*-value=2.2 × 10^-16^); **Figure 5B**. Among the top pathways in MCI (and down-regulated under the music stimuli), are the up-regulated “epigenetic maintenance of chromatin in transcription competent conformation” and “negative regulation of glial cell proliferation”. Additionally, the down-regulated processes included “peptidyl arginine methylation”, “positive regulation of monocyte chemotasis”, “regulation of DNA directed DNA polymerase activity”, “regulation of translational elongation”, and “response to cobalt ion” **(Figure 5B, Table S4)**.

We additionally examined the offset effect of the musical stimuli on the expression of pathways involved in AD, by comparing DEPs_AD_ with musically-stimulated DEPs (**Figure 5C, Table S4**). In 79.1% (174/220) of the FDR DEPs_AD_, log_2_FC values were negatively correlated (ρ=-0.88, *P*-value=2.2 × 10^-16^); this trend dominates the overall correlation value for all shared DEPs (*n*=220; ρ=-0.62, *P*-value=2.2 × 10^-16^); **Figure 5C**. Among the top pathways that are up-regulated in AD, and therefore down-regulated after listening to music in ACD, there are processes involved in DNA repair and replication, such as “mismatch repair”, “regulation of helicase activity” and “regulation of DNA directed DNA polymerase activity”, and others related to protein translation, namely “ribosome disassembly” **(Figure 5C, Table S4)**. Interestingly, the only top pathway that is over-activated in AD (*P*-value=1.4 × 10^-5^) and under-activated after musical stimuli in ACD patients (*P*-value=5.1 × 10^-5^) was the “negative regulation of glial cell proliferation”.

The FDR DEGs_MCI_ and DEPs_MCI_ (*n*=207 and *n*=227) as well as FDR DEGs_AD_ and DEPs_AD_ (*n*=131 and *n*=220, respectively) observed in the comparisons above, overlap in 114 DEGs and 147 DEPs; both genes and pathways also have highly correlated log_2_FC values (ρ [DEGs]=0.88 and ρ [DEPs]=0.94; **Figure S7**). This probably reflects the fact that MCI is often an early or transitional stage between aging and AD [21].

## Discussion

Music exerts a profound influence on human brains, affecting emotions, activating the reward system, and inducing changes in brain anatomy. However, research on the impact of music on gene expression is still in its nascent stages. The present study represents the first attempt to analyze the effect of music on the transcriptome of ACD patients. Capillary blood samples were collected from ACD patients and healthy donors using a minimally invasive procedure before and after a 50-minute session of classical musical stimuli in an ecologically valid environment. Subsequently, we performed RNA sequencing on these samples, allowing for the analysis of gene expression information for over 35,000 transcripts. The analysis focused on differential expression between the two time points and cohorts, as well as the contextualization of gene expression changes within the framework of genes and pathways that exhibit dysregulated expression in MCI and AD (**Figure 6**).

**Figure 6.**
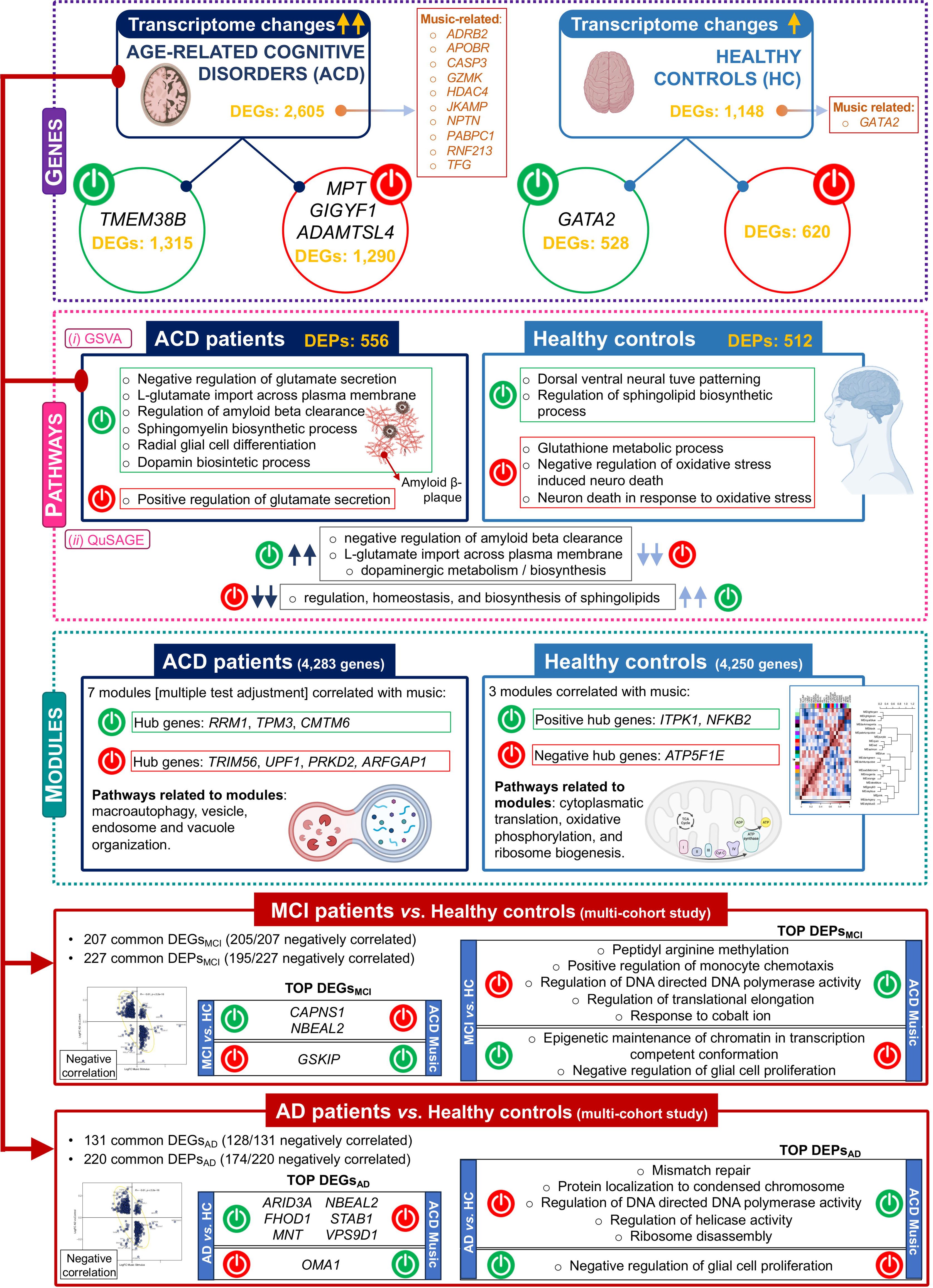
Summary of the most outstanding features related to the impact of musical stimuli on gene expression in the studied cohorts. The figure was built using Biorender resources (https://biorender.com/).

The findings of the present study revealed several noteworthy observations. Firstly, the number of DEGs is more than double in ACD patients compared to healthy controls. Secondly, on average, the transcriptomes of ACD patients quantitatively deviate to a greater extent before and after the music stimuli compared to controls. Additionally, there was a trend towards up-regulation of DEGs in ACD patients, while healthy donors exhibited the opposite effect. These global gene expression changes warrant further investigation, particularly considering the potential massive impact of music’s effect on transcriptomes, which may surpass previous conceptions. Notably, neurologists have posited that music affects almost all regions of the brain [22]; in parallel, our study indicated a high impact of music on our transcriptomes. It would be of particular interest to explore the extent to which this effect underlies the widely held belief that music can help manage AD and other medical conditions [1].

Several genes displayed extreme DE when comparing time points. *GIGYF1*, which is associated with autism spectrum disorder (ASD) and indirectly linked to AD, as well as *ADAMTSL4*, involved in the metabolism of amyloidogenic peptides in AD, were both down-regulated in ACD patients. In contrast, *GATA2*, previously reported to be associated with music [7, 23], was up-regulated in controls; **Supplementary Text**. Pathway analysis revealed important insight into the biological processes impacted by music stimuli. The metabolism of L-glutamate was found to be activated in ACD compared to controls; this pathway is known to play a role in the pathogenesis of AD at early stages [24]. The data suggest that musical stimuli could contribute to managing L-glutamate in the brains of AD patients, which aligns with traditional pharmacological therapies targeting the same pathway [25]. Furthermore, DE of sphingolipid metabolism was observed between ACD and healthy individuals; these complex molecules have been found to be associated with neurodegenerative processes [26]. Music stimuli also impact the regulation of the Aβ peptides in ACD patients. Accumulation of Aβ in the brain is an early toxic event in AD pathogenesis; our study showed that music stimuli targeted the regulation of Aβ with FC more than 15 times greater in ACD patients compared to controls; **Supplementary Text**. Moreover, several articles have found a link between dopaminergic pathways, memory, and music [27, 28]; our results are in line with these previous findings showing that dopamine-related pathways are activated in ACD patients after the musical stimuli.

Analysis of co-expression networks identified clusters of correlated genes showing significant patterns of global activation (positively correlated) or inhibition (negatively correlated) after listening to music. In agreement with analysis of individual genes, musical stimuli target important pathways involved in the pathogenesis of MCI and AD. Functional analysis of the genes in these modules (especially the ‘*TPM3*’ and ‘*CMTM6*’ modules) in ACD patients pointed out the music-mediated activation of biological processes including macroautophagy, vesicle transport, vacuole, and autophagosome organization. Autophagy is a key mechanism to maintain cellular homeostasis by facilitating the clearance of long-lived proteins, including toxic protein aggregates [29], and is essential for maintaining a proper axonal homeostasis [30].

The present study provides compelling evidence indicating that musical stimuli in ACD patients elicit the expression of a notable number of genes and pathways in a direction contrary to that observed in individuals with MCA and AD conditions (**Figure 4**). Several genes stand out as expressing in an opposite direction, including: (1) *CALPNS1* (Calpain Small Subunit 1), which encodes a protein crucial in various biological processes, namely: proliferation, migration, and autophagy, and has been implicated in neurodegenerative processes [31]; (2) *NBEAL2* (Neurobeachin Like 2), previously identified as DE in the brains of individuals with AD [32]; (3) *GSKIP* (GSK3B interacting protein), known to play a role in Tau phosphorylation, a key protein involved in the pathogenesis of AD; (4) *STAB1* (stabilin 1), associated with anxiety [33] and impaired cognition [34]; (5) *OMA1* (OMA1 Zinc Metallopeptidase), which encodes a protease that plays a critical role in maintaining mitochondrial homeostasis [35, 36] (also, impaired processing of *OPA1*, one of the main substrates of *OMA1*, has been reported in post-mortem brains of AD patients [37, 38]); and (6) *FHOD1* (Formin Homology 2 Domain Containing 1), a gene involved in the regulation of actin cytoskeleton organization or gene transcription, and found to be over-expressed in extracellular vesicles of AD patients [39]. An enrichment map of these highly negatively correlated genes reveals their involvement in MCI- and AD-associated pathways, particularly the “regulation of macroautophagy” (when comparing MCI patients to controls) and the “NRAGE cell death” (when comparing AD patients to controls); **Figure S6.** The “receptor of advanced glycation end-products” (RAGE) pathway, including its N-truncated isoform NRAGE, has been implicated in neurotoxic and inflammatory cascades of neurodegenerative processes [40, 41]; **Supplementary Text**.

The impact of music on gene expression related to MCI and AD reveals also a significant compensatory effect when examining DEPs. Notably, several crucial pathways involved in DNA repair/damage response and protein synthesis exhibited remarkable involvement. Previous studies have linked deficiencies in DNA protective mechanisms to age-related neurodegenerative disorders such as Parkinson’s disease and AD [42-44]. Down-regulation of genes involved in the DNA repairing system has been associated with AD [45-47]. Similarly, adequate protein synthesis plays a crucial role in establishing and maintaining long-term memory, while dysfunction in protein homeostasis, synthesis, and ribosomes represents characteristic features of AD [48, 49]. Among ACD patients, music-induced down-regulation was observed in the glial cell proliferation pathway. Glial cells, encompassing microglia, astrocytes, and oligodendrocytes, are non-neuronal cells in the central/ peripheral nervous system (CNS/PNS), crucial for supporting neuronal functions, providing insulation, participating in immune responses, and promoting homeostasis. Dysregulation of glial cells has been associated with AD; however, their specific roles remain unclear [50]. Some researchers suggest that increased activation and proliferation of microglia and astrocytes have neuroprotective effects in AD [51, 52], facilitating Aβ clearance [53-56] and restricting plaque growth [57-59]. Conversely, others propose that microgliosis or astrogliosis processes contribute to AD pathogenesis [60-64]. Additionally, the involvement of oligodendrocytes has been investigated due to their role in myelin production and maintenance [65]. Neuronal demyelination, a hallmark of AD, suggests a potential role for oligodendrocyte loss in this pathological feature [66, 67].

There are several limitations to this study. Firstly, although the study’s population-based nature aimed to minimize noise originating from stimuli other than music, the experimental design is susceptible to potential confounding effects that may be unavoidable. Secondly, our findings need to be investigated and validated in other ecological settings. Thirdly, the applied sampling strategy does not allow an investigation into the long-term effects of music on gene expression, or its potential connections to the benefits attributed to music in patients with MCI and AD; **Supplementary Text**.

Genes and pathways related to the central nervous system (CNS) exhibit the highest sensitivity to musical stimuli, which is expected given that sight and hearing serve as natural entry points for such stimuli. Furthermore, the transcriptomes of ACD patients display greater sensitivity to music and undergo more significant changes compared to healthy individuals. The observation that many genes/pathways influenced by music exhibit an opposite direction of expression and regulation in comparison to those altered in MCI/AD warrants careful attention in future studies, particularly in exploring the potential therapeutic implications of music for neurodegenerative diseases. Validating the current findings in larger and independent cohorts is imperative. Additionally, expanding the study to include other disorders i.e., mental-related disorders, TEA, brain damage, etc, would provide a broader understanding of the effects of music on gene expression. Further research is necessary to fully comprehend the implications of these findings for human health [1].

## Ethics approval and Consent to participate

Written informed consent was obtained from all the participants in the present study. The Ethics Committee of Xunta de Galicia approved the present project (Registration code: 2020/021), and the study was conducted in accordance with guidelines of the Helsinki Declaration.

## Consent for publication

All participants have given permission for publication of the project’s findings.

## Availability of data and materials

The authors confirm that data supporting the findings of this study are available in Gene Expression Omnibus – NCBI (GEO; https://www.ncbi.nlm.nih.gov/geo/) under accession number XXXX.

## Author’s contribution

AS, FM-T and LN conceived and coordinated the study; AS and AGC analyzed the data and wrote the first draft; JP-S, XB, SP, SV-L, AC-M, M-JC, IF, N;, SR-V, LR, AD-U, FC-V, IR-C, CR-T, FM-T, AS, contributed to logistics and sample and data collection; All the authors revised and approved the final version of the text.

## Acknowledgements

We would like to kindly acknowledge all participants in the present study, the AGADEA association of Alzheimer disease patients, and the musicians of SANARTE, who kindly agreed to participate in this project. We would also like to acknowledge the Real Filharmonía de Galicia (www.rfgalicia.org; and Sabela García Fonte in particular), and the Auditorio de Galicia for their support. This study received support from GAIN (IN607B 2020/08 [A.S.] and IIN607A2021/05 [F.M-T]).

We would like to thank the contribution of all members of the Sensogenomics

Working Group:

Antonio Salas Ellacuriaga – PI

Federico Martinón-Torres – PI;

Laura Navarro Ramón – Coordinator

## GenPoB/GenVip - Instituto de Investigación Sanitaria (IDIS) (alphabetic order)

Alba Camino Mera, Albert Padín Villar, Alberto Gómez Carballa, Alejandro Pérez López, Alicia Carballal Fernández, Ana Cotovad Bellas, Ana Isabel Dacosta Urbieta, Narmeen Mallah, Ana María Pastoriza Mourelle, Ana María Senín Ferreiro, Andrés Muy Pérez, Antía Rivas Oural, Antonio Justicia Grande, Antonio Piñeiro García, Anxela Cristina Delgado García, Belén Mosquera Pérez, Blanca Díaz Esteban, Carlos Durán Suárez, Carmen Curros Novo, Carmen Gómez Vieites, Carmen Rodríguez-Tenreiro Sánchez, Celia Varela Pájaro, Claudia Navarro Gonzalo, Cristina Serén Trasorras, Cristina Talavero González, Einés Monteagudo Vilavedra, Estefanía Rey Campos, Esther Montero Campos, Fernando Álvez González, Fernando Caamaño Viñas, Francisco García Iglesias, Gloria Viz Rodríguez, Hugo Alberto Tovar Velasco, Irene Álvarez Rodríguez, Irene García Zuazola, Irene Rivero Calle, Iria Afonso Carrasco, Isabel Ferreirós Vidal, Isabel Lista García, Isabel Rego Lijo, Iván Prieto Gómez, Iván Quintana Cepedal, Jacobo Pardo Seco, Jesús Eirís Puñal, José Gómez Rial, José Manuel Fernández García, José María Martinón Martínez, Julia Cela Mosquera, Julia García Currás, Julián Montoto Louzao, Lara Martínez Martínez, Laura Navarro Marrón, Lidia Piñeiro Rodríguez, Lorenzo Redondo Collazo, Lúa Castelo Martínez, Lucía Company Arciniegas, Luis Crego Rodríguez, Luisa García Vicente, Manuel Vázquez Donsión, María Dolores Martínez García, María Elena Gamborino Caramés, María Elena Sobrino Fernández, María José Currás Tuala, María Martínez Leis, María Soledad Vilas Iglesias, María Sol Rodriguez Calvo, María Teresa Autran García, Marina Casas Pérez, Marta Aldonza Torres, Marta Bouzón Alejandro, Marta Lendoiro Fuentes, Miriam Ben García, Miriam Cebey López, Montserrat López Franco, Narmeen Mallah, Natalia García Sánchez, Natalia Vieito Perez, Patricia Regueiro Casuso, Ricardo Suárez Camacho, Rita García Fernández, Rita Varela Estévez, Rosaura Picáns Leis, Ruth Barral Arca, Sandra Carnota Antonio, Sandra Viz Lasheras, Sara Pischedda, Sara Rey Vázquez, Sonia Marcos Alonso, Sonia Serén Fernández, Susana Rey García, Vanesa Álvarez Iglesias, Victoria Redondo Cervantes, Vanesa Álvarez Iglesias, Wiktor Dominik Nowak, Xabier Bello Paderne, Xabier Mazaira López

## Nursing volunteers (alphabetic order)

Alejandra Fernández Méndez, Ana Isabel Abadín Campaña, Ana María León Caamaño, Ana María Buide Illobre, Ángeles Mera Cores, Carmen Nieves Vastro, Carolina Suarez Crego, Concepción Rey Iglesias, Cristina Candal Regueira, Dolores Barreiro Puente, Elvira Rodríguez Rodríguez, Eugenia González Budiño, Eva Rey Álvarez, Fernando Rodríguez Gerpe, Gemma Albela Silva, Isabel Castro Pérez, Isabel Domínguez Ríos, José Ángel Fernández de la Iglesia, José Cruces Vázquez, José Luis Cambeiro Quintela, José Ramón Magariños Iglesias, Julia Rey Brandariz, Julio Abel Fernández López, Luisa García Vicente, Manuel González Lito, Manuel González Lijó, Manuela Pérez Rivas, Margarita Turnes Paredes, María Aurora Méndez López, María Begoña Tomé Arufe, María Campos Torres, María del Carmen Baloira Nogueira, María del Carmen García juan, María Esther Moricosa García, María Luz Chao Jarel, María Martínez Leis, María Mercedes Jiménez Santos, María Salomé Buide Illobre, María Victoria López Pereira, Mercedes Jorge González, Mercedes Isolina Rodríguez Rodríguez, Miren Payo Puente, Natalia Carter Domínguez, Olga María Reyes González, Pilar Mera Rodríguez, Purificación Sebio Brandariz, Salomé Quintáns lago, Yolanda Rodríguez Taboada, María Pereira Grau.

## Other volunteers (alphabetic order)

Alba Arias Gómez, Alejandro Moreno Díaz, Ana Arca Marán, Astro González Guirado, Brais García Iglesias, Carlos Sánchez Rubín, Carmen Otero de Andrés, Clara Pérez Errazquin Barrera, Claudia Rey Posse, Cristina Rojas García, Eduardo Xavier Giménez Bargiela, Elena Gloria Morales García, Fabio Izquierdo García Escribano, Gabriel Guisande García, Jaime López Martín, Lara Pais Ramiro, Lucía Rico Montero, Luís Estévez Martínez, Manuel Estévez Casal, María Aránzazu Palomino Caño, María Rubio Valdés, Marisol Nogales Benítez, Miryam Tilve Pérez, Nuria Villar Muiños, Pablo Del Cerro Rodríguez, Pablo Pozuelo Martínez Cardeñoso, Salma Ouahabi El Ouahabi, Santiago Vázquez Calvache

## Supplementary Data

**Table S1.** DEGs detected between TP1 and TP2 in the ACD and healthy control cohorts.

**Table S2.** Biological processed terms differentially activated in ACD and healthy controls between TP1 and TP2, as derived from GSVA analysis; and neuro-biological related terms differentially activated between ACD patients and healthy controls in response to musical stimuli, as derived from QuSAGE analysis.

**Table S3.** Modules of co-expressed genes detected in the AD patients and healthy controls cohorts and their correlation with the musical stimuli (FDR=false discovery rate). Over-representation analysis of ACD and healthy controls modules, using Gene Ontology (GO) database as reference.

**Table S4.** Correlation between shared DEGs and DEPs obtained from MCI/AD *vs*. controls (multi-cohort case-control study) and from TP2 *vs*. TP1 in ACD patients (Pretest-Posttest study). Enrichment analysis results from the PPI network of the shared DEGs. In the enrichment map, nodes represent gene-sets and edges represent mutual overlap (highly redundant gene-sets are grouped together as clusters) [12].

**Figure S1.** Expression values heatmap of all DEGs (*P*-value <0.05) between TP1 and TP2 in both ACD and healthy controls cohorts.

**Figure S2.** GSVA showing differences detected in the activation or inhibition of the main neuro-biological pathways in patients and controls when exposed to music (FDF<0.05)

**Figure S3.** Correlation between activation values (represented as log_2_FC) from shared significant (FDR<0.05) neuro-biological pathways detected in ACD and healthy controls cohorts.

**Figure S4.** QuSAGE analysis between ACD patients and healthy controls. (A) QuSAGE results showing the top differentially activated pathways between ACD patients and controls related to neuro-biological processes (*P*-value<0.05; |*PA_ACD_*|+|*PA_HC_*|>0.05). (B) Activity of individual genes belonging to the pathway “negative regulation of amyloid beta clearance”.

**Figure S5.** Co-expression network analysis in ACD patient’s cohort. A) Clustering dendrogram of genes and co-expression modules detected, represented by different colors. (B) Hierarchical clustering eigengene dendrogram and heatmap for ACD patients’ datasets showing relationships among the modules and TP (musical stimuli; TP2 *vs.* TP1). Gene names on the left of the heatmap are the hub genes of each module. (C) Plots showing comparison between MM (module membership) and musical stimuli correlation of genes from the significant (FDR<0.05) modules detected in the ACD cohort. **Figure S6.** PPI network and downstream enrichment analysis of the proteins coded by the top A) 207 DEGs in MCI *vs*. healthy controls, and B) 131 DEGs in AD *vs*. healthy controls, and impacted by musical stimuli in ACD patients (FDR < 0.05 in both studies).

**Figure S7.** Correlation of log_2_FC for the DEGs and DEPs detected in MCI (when compared against healthy controls) and the DEGs and DEPs detected in AD (when compared against healthy controls) that are also significant in ACD after the musical stimuli (see **Table S4**, **Figure 4 and Figure 5** for details). For all the correlation values, *P*-value < 2.22×10^-16^.

