## Supplementary figures and images for "Tuning gene expression to music: the compensatory effect of music on age-related cognitive disorders"

### Figure S1.tif

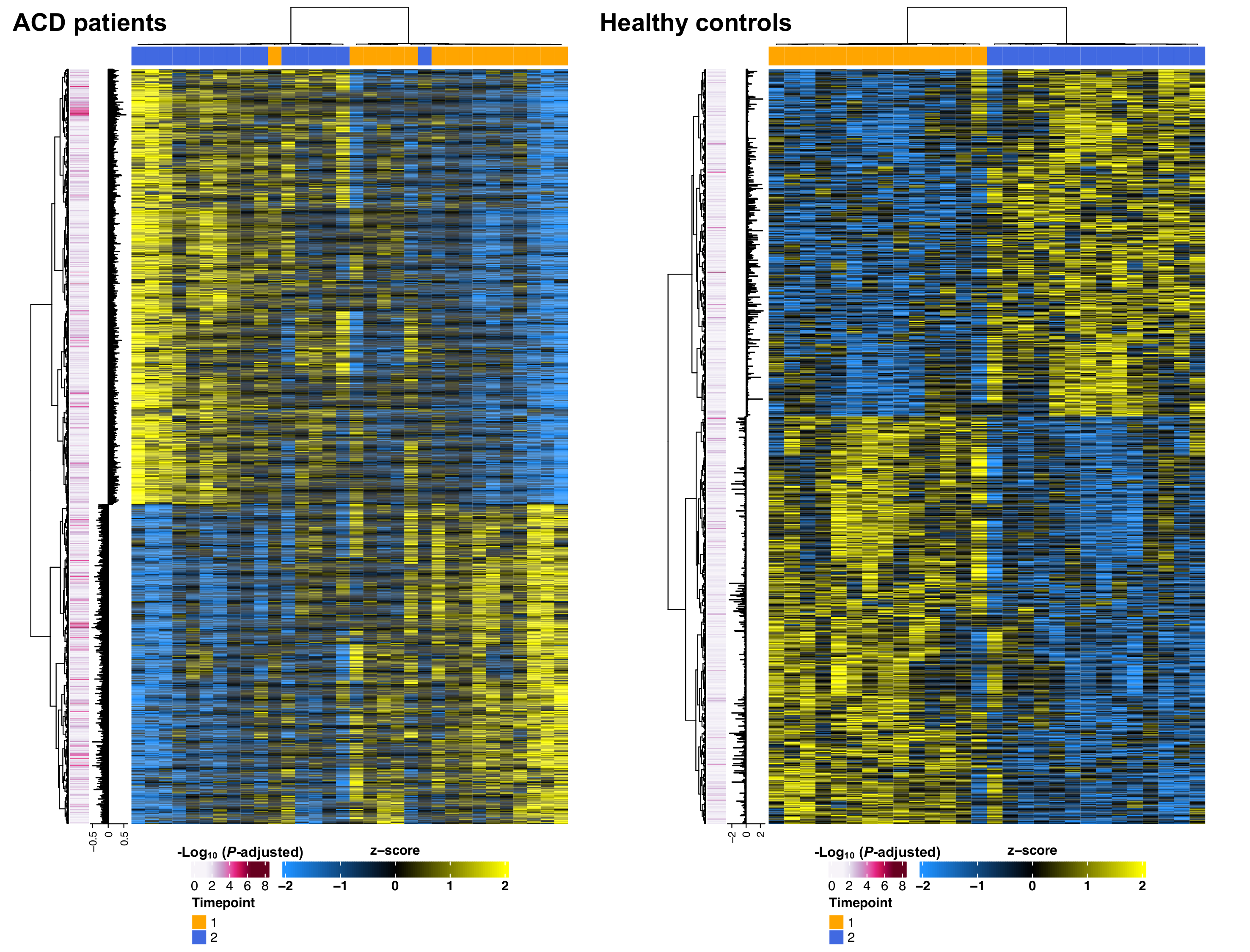

### Figure S2.tif

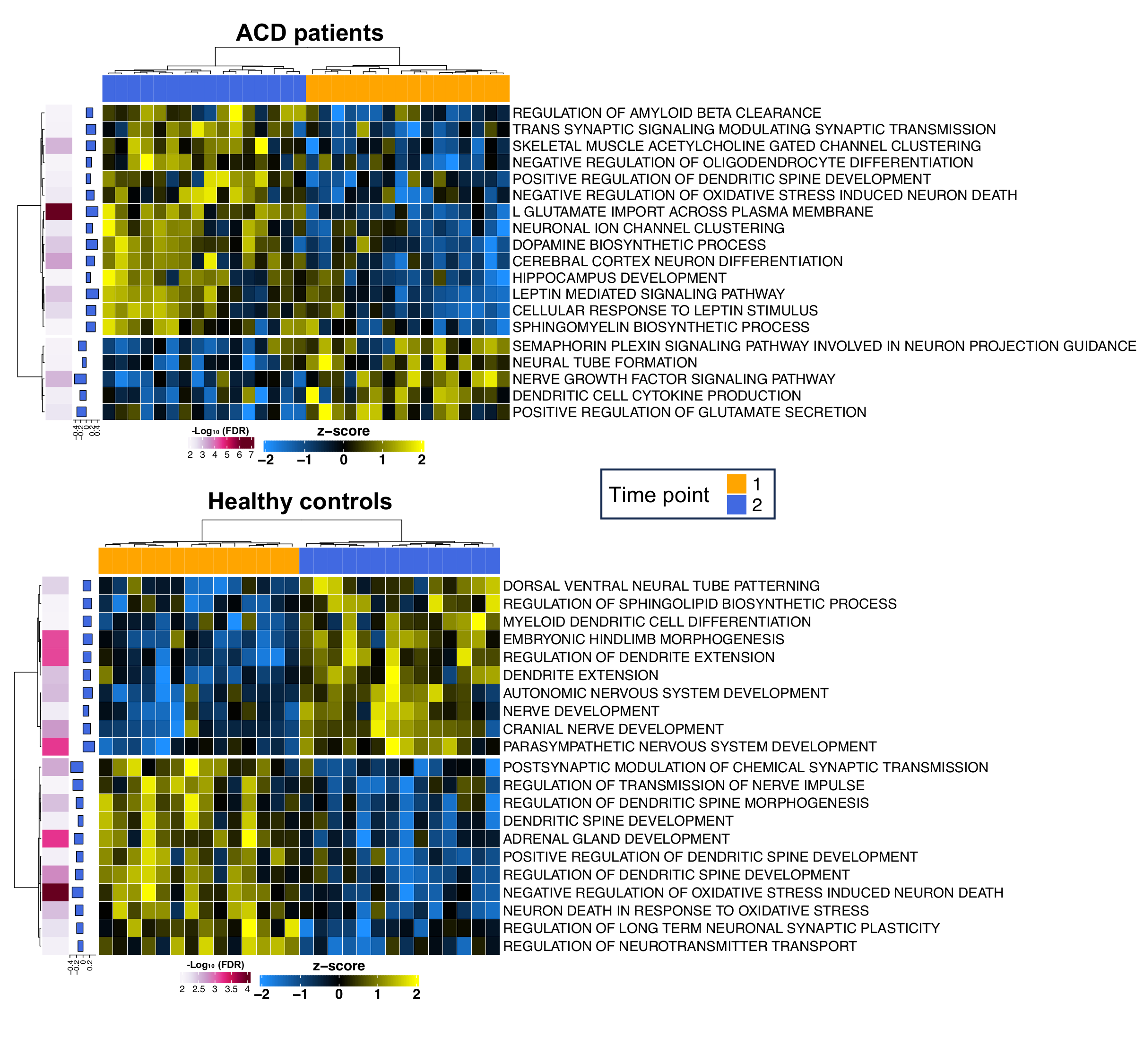

### Figure S3.tif

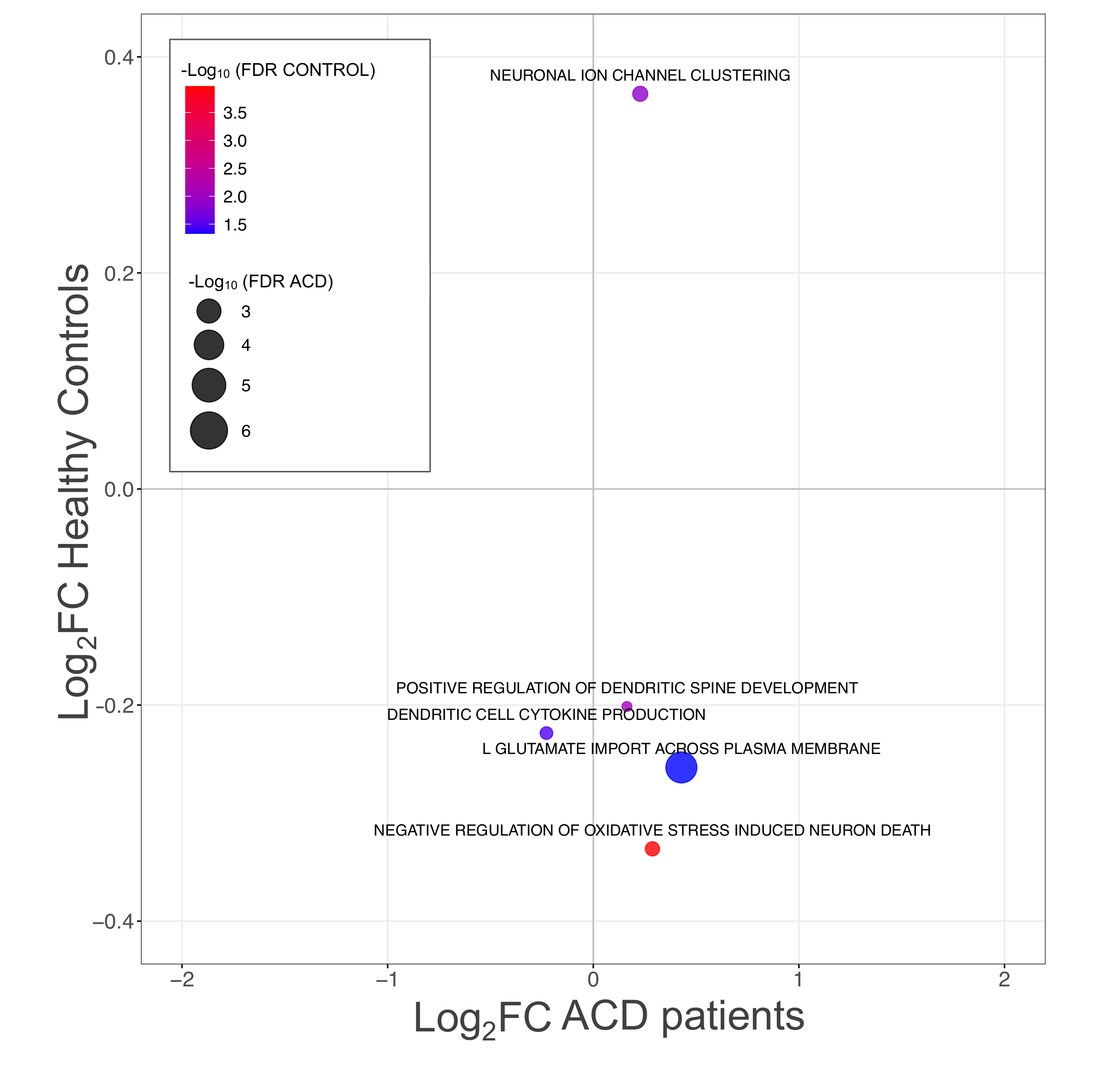

### Figure S4.tif

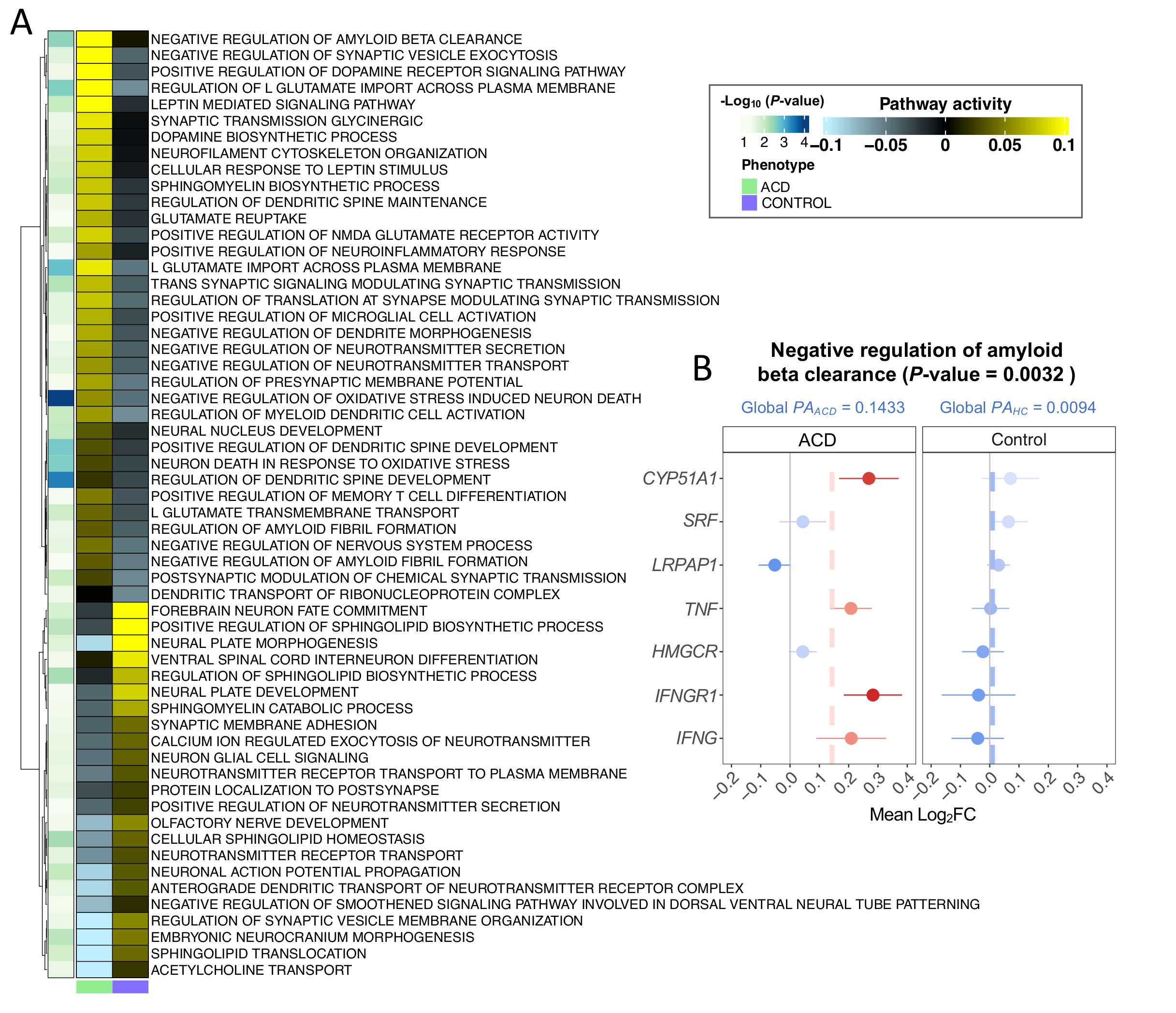

### Figure S5.tif

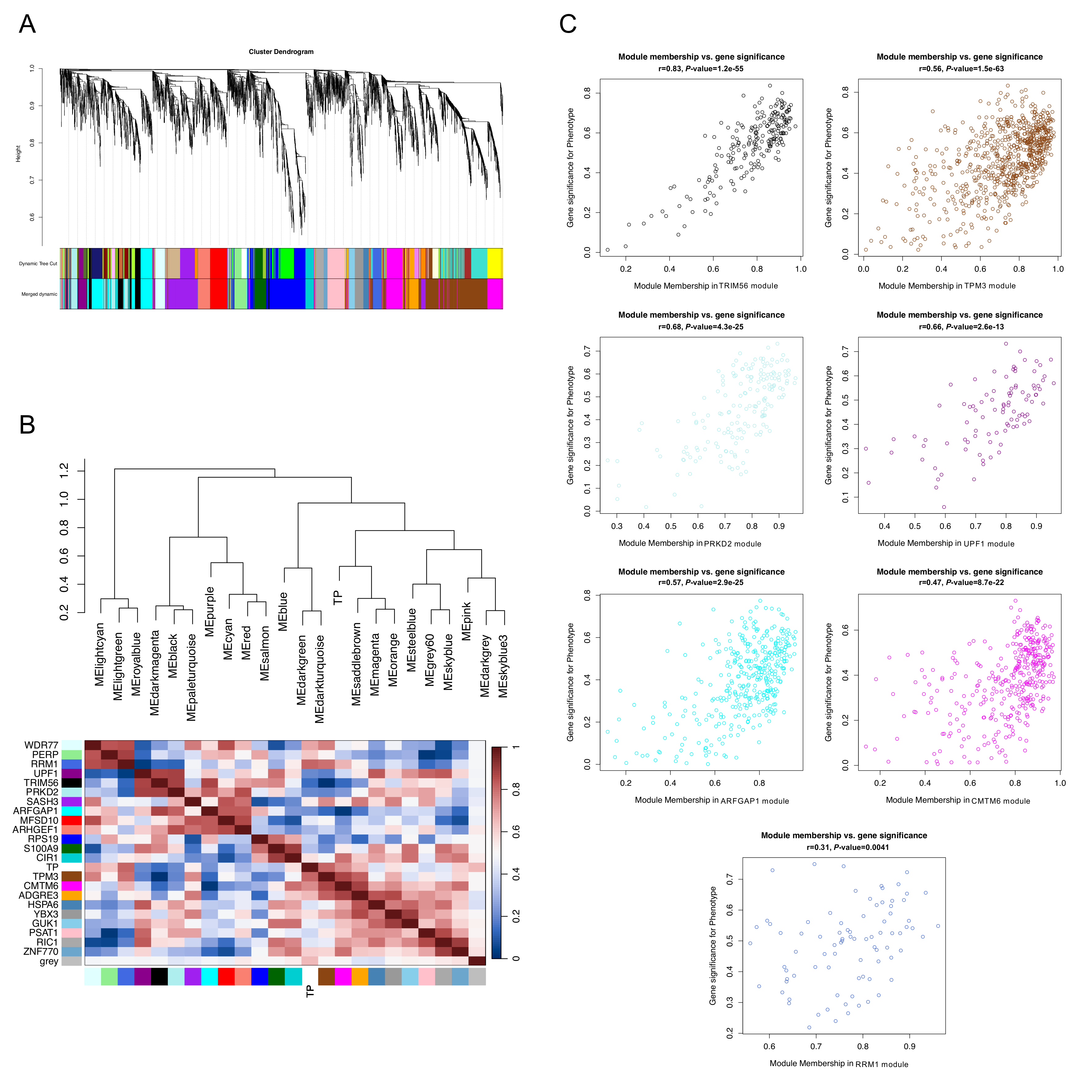

### Figure S6.tif

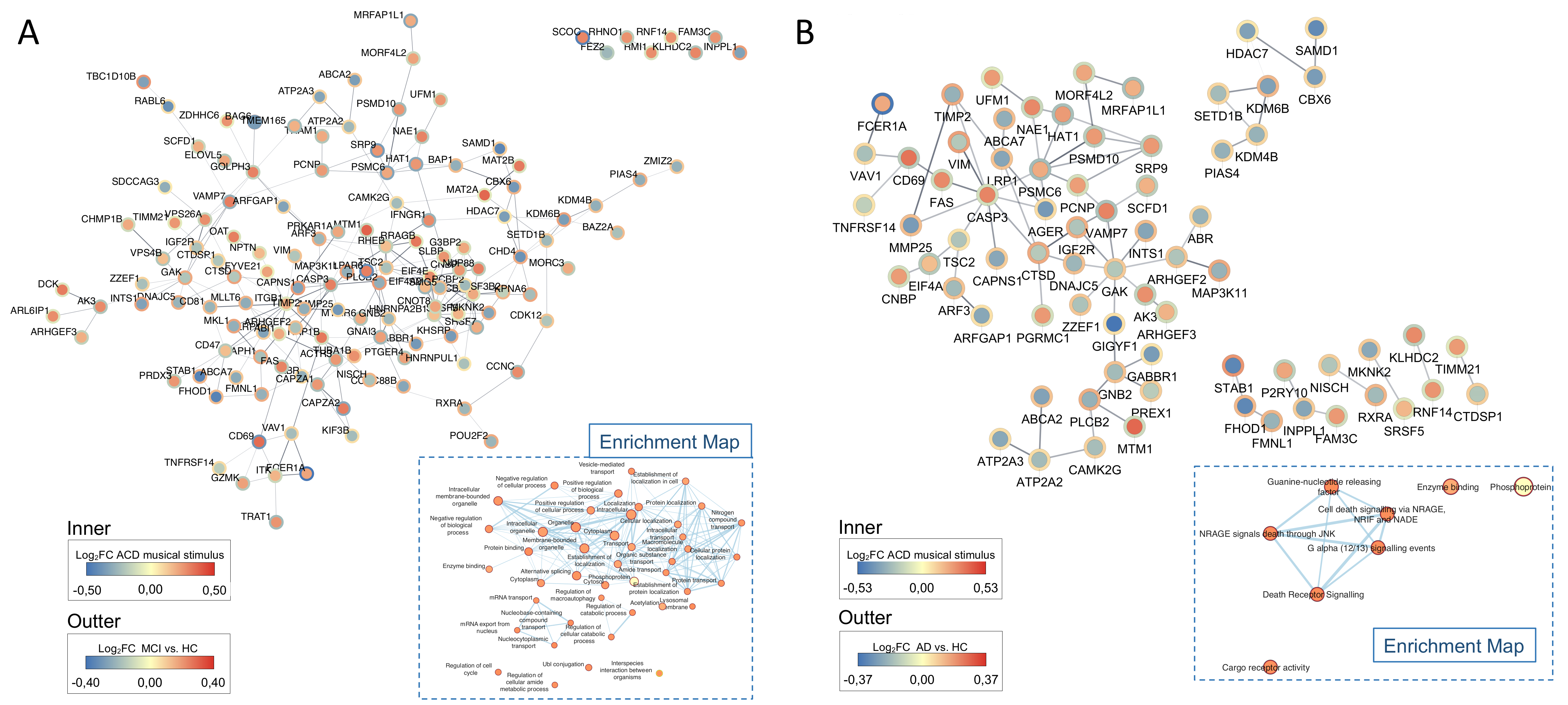

### Figure S7.tif

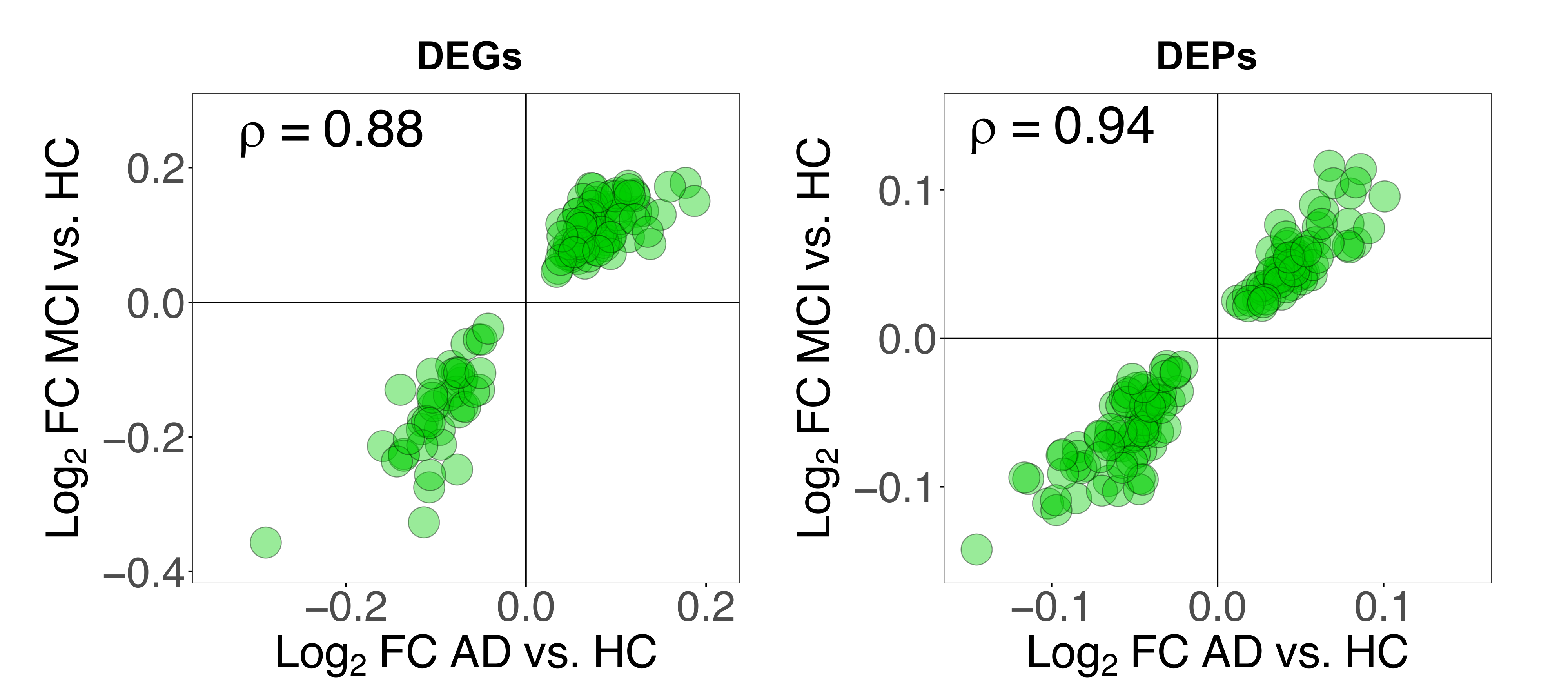
