## Supplementary Text for "Tuning gene expression to music: the compensatory effect of music on age-related cognitive disorders"

**Additional background**

Santiago Ramón y Cajal, who was awarded the Nobel Prize in Physiology and Medicine in 1906, and his colleagues affirmed that “*...the ability of a pianist … requires many years of mental and muscular gymnastics. To understand this important phenomenon, it is necessary to accept that … new pathways are created by the ramification and progressive growth of terminal dendritic and axonal processes*” [1]; Wan and Schlaug [2] have also quoted this statement. In the early days of research, the claim that music can induce changes in the brain’s anatomy was supported by limited and indirect evidence. However, current scientific evidence amply supports this proposition. For instance, Bermudez et al. [3] demonstrated that the thickness of the cortex, with peaks in the superior temporal and dorsolateral frontal regions, is greater in musicians compared to non-musician controls. In a review conducted by Wan and Schlaug [2], the existing evidence of music-making as a tool for promoting brain plasticity was analyzed, concluding that the human brain can be shaped by musical experiences. The review also pointed out the beneficial use of music as a non-pharmacological therapy in neurological and developmental disorders.

Furthermore, Fischer et al. [4] indicated that repeated exposure to personally meaningful music stimulates neural connectivity, leading to improvements in cognitive functioning in patients with early-stage cognitive decline. This finding suggests that music may have therapeutic applications in patients with dementia.

In addition, the emotional power of music has been a topic of interest since classical Greece, and especially during the Baroque period, where the doctrine of affections explored the relationship between music and emotions. Presently, there is scientific support for the neurological mechanisms underlying the ability of music to evoke powerful emotional responses [5]. Pereira et al. [6] used functional magnetic resonance imaging (fMRI) techniques to demonstrate that emotion-related limbic and paralimbic regions, as well as the reward circuitry, were more active when participants listened to familiar music compared to unfamiliar music. The study by Logeswaran and Bhattacharya [7] demonstrated that the emotions of music are cross-modal, such that they can spread from one sensory system to another. Additionally, using fMRI, Salimpoor et al. [8] demonstrated that neuronal activity in the mesolimbic striatum, particularly the nucleus accumbens, during the listening of a novel, unheard piece of music was the best predictor of the amount of money listeners were willing to spend on purchasing the piece. This study also highlighted the involvement of sensory cortical areas in reward processing.

In a recent systematic review on neuroscience and cognitive sciences, we [9] highlighted the existence of a global consensus on the beneficial effects of music on human health, particularly in relation to Alzheimer’s Disease (AD). According to this review, more than 93% of neuroscientific studies focused on music and neurodegenerative disease indicated a positive contribution of music to human health. The benefits of music were primarily associated with memory and cognition, as well as social behavior, mood, and emotion. This conclusion aligns with those of the World Health Organization (WHO). In a scoping review on health and well-being, the WHO recognized the growing evidence base for the role of the arts, including music, in improving health and well-being (https://www.who.int/europe/publications/i/item/9789289054553): *“the growing evidence base for the role of the arts in improving health and well-being by e.g. supporting the implementation of arts interventions where a substantial evidence base exists, such as the use of recorded music for patients prior to surgery, arts for patients with dementia and community arts programmes for mental health*”.

**Additional information on methods**

Participants, RNA isolation and RNA-Seq analysis

The experimental concert took place in the Auditorio de Galicia, which is renowned both nationally and internationally for hosting important classical music events. The participants in this study consisted on two cohorts: the first cohort comprised members of the ‘Asociación Galega de Axuda ós Enfermos con Demencias tipo Alzheimer’ (AGADEA) association, who suffer from age-related cognitive disorder (ACD). The second cohort served as the healthy control group and consisted of patient’s caregivers, and musicians. The study ensured that participants experienced a comfortable and welcoming environment during the classical music concert, minimizing exposure to stress factors that could potentially influence the experiment and bias the results. Several weeks prior to the Sensogenomics22-pilot concert, the project`s organizing team conducted informative sessions with donors at the mentioned auditorium, as well as at the Alzheimer’s Association offices and centers. AGADEA offers various local programs, resources, and services to patients, including home care, adult day care, and meal services. These sessions provided an opportunity to obtain written informed consent for sample collection. Additionally, a questionnaire was administered to collect data on the donors’ pathological conditions and self-reported music education. Organizing these sessions well in advance of the experimental concert allowed for the prevention and minimization of potential stress factors on the day of the musical event, which could have negatively impacted the donors’ transcriptomes.

The musical repertoire lasted 50 minutes and included short pieces of classical music interpreted by different combinations of an initial septet (the SANARTE musical group, made of two violins, two violas, two cellos, and one horn) conformed by professional musicians (some were members of the Real Filharmonía de Galicia or RFG; www.rfgalicia.org). The repertoire consisted of adaptations of the “Spring” from the “Four Seasons” by Antonio Vivaldi, “A little night music, Allegro” by Wolfgang Amadeus Mozart, Pavane pour une Infante Defunte by Maurice Ravel, “La musica notturna delle strade di Madrid” by Luigi Boccherini, “Rondo in E-flat major, K.371” by Wolfgang Amadeus Mozart, the “Andante for Horn and Piano” by Richard Strauss, and “Por una cabeza” by Carlos Gardel. To prevent the audience (donors) from being influenced by the style and specific pieces of music being performed, the repertoire was not disclosed beforehand.

A total of 100–200 μl of blood was taken from all the donors using a capillary puncture procedure. The sample was first collected using a Minivette^®^ POCT K3 EDTA device and immediately transferred to microtubes containing an appropriate volume of PAXgene additive to stabilize the blood and preserve RNA integrity. On average, the sample collection process lasted approximately 2–3 minutes for each donor, and all the samples were collected within two short time frames (less than 30 minutes) immediately before and after the concert.

RNA isolation and RNA-Seq analysis

Total RNA was isolated using the PAXgene blood miRNA extraction kit (Qiagen) adapting buffer volumes to the amount of blood collected (100–200μl). On-column DNase I treatment was carried out during the extraction process. RNA concentration step and an additional DNase treatment were undertaken using an RNA clean & concentrator kit (Zymo Research) prior to sequencing. The amount and integrity of the RNA obtained were checked using the TapeStation 4200 (Agilent). Input material was normalized, and strand specific library preparation was completed using the SMARTer Ultra low input RNA library (Poly-A). Sequencing was performed using a NovaSeq 6000 System at 100 paired end configuration to finally obtain 70M reads per sample. Quality control (QC) of the RNA-Seq data obtained was performed using the *FastQC* and *MultiQC* [10]. After the QC step, problematic 3' end and adapters were removed with the *Trimmomatic* package [11], and the reads mapped against the human reference genome (GCh38,v.104) using ultrafast universal RNA-Seq aligner *STAR* [12]. Secondary and low-quality alignments were removed using *SAMtools* [13]. To count the number of reads mapping to each gene (counts per gene), we used the software *HTSeq-counts* [14].

Pathway analysis

Biological pathways involved in musical stimulation were inferred directly from gene expression data following two different approaches. First, we used normalized data (corrected for paired sampling) as input in the Gene Set Variation Analysis (*GSVA*) R package [15] to detect biological processes that significantly change between baseline and TP2 in each of the groups separately (ACD patients and controls). We used the *gsva* method with a minimum gene-set size of 10 to estimate the gene-set enrichment scores per sample. Differential pathway expression analysis was carried out with *limma* [16]. This procedure was also employed to detect differentially expressed pathways (DEPs) in the AD case-control multi-cohort study. Second, two-way pathway analysis was performed using the Quantitative Set Analysis for Gene Expression (*QuSAGE*) R package [17] to find specific differential responses in biological processes to the musical stimuli between ACD patients and the control group. *QuSAGE* quantifies gene set activity using a complete Probability Density Function (PDF). As in the case of individual gene approach, we accounted for patient specific effects to calculate pathway activation by including sample information in the design formula. In both *GSVA* and *QuSAGE* analyses, we used the ontology gene-sets C5 (biological processes subset) as the reference database, contained in the file *gmt*, downloaded from the human Molecular Signatures DataBase (MSigDB) [18]. For some specific analysis, we used a sub-selection of neuro-biological processes by systematically and manually curating the pathway analysis results for items containing a neuro-biological term; this selection yielded a total of 483 biological processes (**Supplementary Text Table 1**).

Co-expression analysis

We investigated clusters of co-expressed genes potentially correlated to the musical stimuli in both ACD patients and the healthy donor group separately. Gene expression data normalized and corrected for patient-to-patient differences was used to build a signed weighted correlation network with the Weighted Gene Co-expression Network Analysis (*WGCNA*) R package [19]. Only genes that showed the most variant expression values between samples (the top 75% with the highest variance) were included in the analysis. A matrix of correlations between all pairs of selected genes was generated from the expression values, and further converted into an adjacency matrix with a power function. We chose a soft-thresholding power based on the criterion of scale-free topology after testing a set of candidate powers. A soft-thresholding power of 20 was selected as it resulted in the maximum model fitting index for both datasets (>0.85; **Supplementary Text Figure 1**). The Topological Overlap Matrix (TOM) and the corresponding dissimilarity (1–TOM) values were calculated. A minimum module size of 30, and 0.2 as the dendrogram cut height threshold for module merging were selected. The detected modules of co-expressed genes were labelled by colors and used to calculate module eigengenes (the PC1 of the module). The correlation between module eigengenes and the clinical trait was analyzed to identify modules of interest significantly associated with the musical stimuli (gene significance, or GS). Module Membership (MM) was calculated as a measure of intramodular connectivity. The hub genes within the significant modules are those with the highest connectivity; we named the modules by their most significant Hub genes.

We studied the biological significance of the correlated modules by performing an over-representation analysis and using the compare cluster function included in the *Clusterprofiler* R package [20] and the biological processes terms from the Gene Ontology (GO) database [21].

**Additional results**

Impact of music stimuli regarding previously reported music-related candidate genes

There is a set of candidate genes (*n* = 334) that have been previously reported to be related to music; their association is mainly inferred from genome association and transcriptomic studies [9]. We cross-compared this set of genes with those that appear as DE when contrasting the two time points. A total of 48 (out of 334; 14.4%; **Supplementary Text Table 2**) music-related genes appear as DEGs in TP2 *vs*. TP1 in ACD patients; among the most significant DEGs (those surpassing the multiple test adjustment) are (**Supplementary Text Figure 2A**): *PABPC1* (*P*-value = 3.5×10^-9^), *ADRB2* (*P*-value = 1.6×10^-4^), *GZMK* (*P*-value = 4.6×10^-7^), *CASP3* (*P*-value = 4.6×10^-5^), *APOBR* (*P*-value = 5.3×10^-6^), *JKAMP* (*P*-value = 1.1×10^-4^), *RNF213* (*P*-value = 6.9×10^-6^), *NPTN* (*P*-value = 9.1×10^-7^), *HDAC4* (*P*-value = 1.1×10^-4^), and *TFG* (*P*-value = 1.9×10^-5^). When examining controls, we detected 25 music-related genes (7.5%) that appear as DEGs in TP1 *vs*. TP2, with *GATA2* (see **Figure 2E** from main text) being the only one surpassing the multiple test adjustment.

DEGs and neuro-biological processes

To better investigate the nature of the DEGs emerging from the TP1 *vs*. TP2 comparison in ACD patients and controls, we examined GO biological terms related to neuro-biological processes and the genes that are involved in these terms. There are 2,192 protein coding genes among the DEGs in ACD patients; of these, 479 fall into neural-related processes, representing 22% of the whole protein coding DEGs (TP1 *vs*. TP2). Ten DEGs in ACD patients falling within these terms showed *P*-values < 10^-5^: *GNPAT* (*P*-value = 6.7×10^-6^), *UFM1* (*P*-value = 9.5×10^-6^), *GCH1* (*P*-value = 1.9×10^-6^) being up-regulated in TP2 with respect to TP1, and *ARHGEF2* (*P*-value = 2.7×10^-6^), *LAMB1* (*P*-value = 8.0×10^-6^), *DNAJC5* (*P*-value = 6.7×10^-6^), *AKT2* (*P*-value = 8.0×10^-6^), *JAK3* (*P*-value = 9.5×10^-6^), *FHOD1* (*P*-value = 8.0×10^-6^), and *SETD5* (*P*-value = 1.5×10^-6^) down-regulated; **Supplementary Text** **Figure 2B**. In healthy controls, there are 176 DEGs (TP1 *vs*. TP2) falling into neural-related processes, representing 19% of the protein coding DEGs (*n* = 921). Five out of the 176 DEGs in these neural categories showed a *P*-value < 10^-5^: two of them were significantly up-regulated (*GATA2* and *CCR5*), and three significantly down-regulated after musical stimuli (*PLCB1*, *ALCAM* and *PRKG1*); **Supplementary Text** **Figure 2B**.

Modules of co-expressed genes associated to musical stimuli in healthy control samples

In the control dataset, 15 modules of co-expressed genes were detected (**Supplementary Text Figure 3A; Supplementary Text Figure 3B**; **Table S6** cited in the main text), 3 of them showing a statistically significant correlation with the musical stimuli (although not significant after multiple test correction). Globally, *ITPK1* [midnight blue] module (R = 0.42, *P*-value = 0.025) and *NFKB2* [greenyellow] module (R = 0.45, *P*-value = 0.016) were positively correlated (**Supplementary Text Figure 3C**), whereas the most significant module in controls, the *ATP5F1E* [pink] module (*P*-value = 0.0083), was negatively correlated with the musical stimuli (R = -0.49; **Table S6** cited in the main text) and, as observed in the module expression patterns of individual samples, values tend to be lower in TP2 with respect to TP1; **Supplementary Text Figure 3D**. The genes most inter-connected within the *ATP5F1E* module also showed a high significant correlation with the musical stimuli (R = 0.38, *P*-value =1.8×10^-7^); **Supplementary Text Figure 3E**. Functional assessment of genes included in the *ATP5F1E* module pointed to processes mainly involved in cytoplasmatic translation (adjusted *P*-value = 3.63×10^-31^), oxidative phosphorylation (adjusted *P*-value = 1.41×10^-11^) and related processes (aerobic respiration and ATP synthesis in the mitochondria), and ribosome biogenesis (adjusted *P*-value = 1.05×10^-5^); **Table S6** cited in the main text; **Supplementary Text Figure 4A**. Although of much less importance, the positively correlated *ITPK1* module could also be linked to activation of T-cell pathways. Genes from the most significant pathways belonging to the *ATP5F1E* module showed high MM values; **Supplementary Text Figure 4B.**

**Additional discussion**

Differential expressed genes in ACD and healthy controls

*GIGYF1* (GRB10-interacting gyf protein 1)

The GRB10-interacting gyf protein 1 (*GIGYF1*) gene is associated with autism spectrum disorder (ASD). Chen et al. [22] indicated that *GIGYF1* deficiency in mice led to a reduction of upper layer cortical neurons accompanied by decreased proliferation and increased differentiation of neural progenitor cells; these authors found that ASD individuals with *GIGYF1* likely gene disruptive (LGD) mutations were less likely to have cognitive impairments. According to these findings, it might be most relevant to investigate if a down-regulated *GIGYF1* (driven in the present study by the musical stimuli) could have some impact on cognitive benefits. Remarkably, *GIGYF1* (together with other genes) is associated with rapamycin sensitive genes (and it has been included in a patented invention for the prevention, treatment, and diagnosis of AD, namely, Therapeutic targets for AD - PCT/GB2O13/051843). In animal models, rapamycin has demonstrated to be the most effective pharmacological treatment to target the aging process to increase life and health span [23], and it also allows to delay/reverse age-related diseases (in mice, rats, and companion dogs), e.g., cancers, cardiac dysfunction, kidney disease, obesity, cognitive decline, periodontal disease, macular degeneration, muscle loss, stem cell function, and immune senescence [24-26]. Aggregation of amyloid beta (Aβ) peptides is a key feature of AD.

*ADAMTSL4*

It has been reported that the DEG (TP1 *vs*. TP2) *ADAMTS4* in ACD encodes for a disintegrin and metalloproteinase with thrombospondin motifs 4 enzyme. This metalloproteinase generates N-truncated Aβ4-x species and marks oligodendrocytes as a source of amyloidogenic peptides in AD [27]. *ADAMTS4* was included in the AD-specific network inferred from genetic association and interactome studies [28]. It was found highly expressed in the hippocampus of children with Autism Spectrum Disorder [29].

*GATA2* (GATA Binding Protein 2)

In healthy controls, the GATA Binding Protein 2 (*GATA2*) was found to be up-regulated in healthy controls when contrasting TP2 *vs*. TP1. This gene encodes a member of the GATA family of zinc-finger transcription factors that regulates GABAergic neurogenesis and migration [30] in adults. It has been discussed several times in the context of music and in the very few transcriptomic studies reported to date [31-35]. Recent studies have identified *GATA2* as regulator of key genes associated with neurodegenerative diseases, such as vascular dementia [36] and AD [37]. It also plays an essential role activating the expression of the neuroglobin gene, which encodes for a protein related to neuroprotection and involved in mitigating the consequences of stroke and AD [38].

Differential expressed pathways in ACD and healthy controls

*Metabolism of L-glutamate*

L-glutamate is the most abundant free amino acid in a healthy brain and the CNS, and it is a major excitatory neurotransmitter in mammals [39] with a main role in neuronal plasticity and synapsis, and also implicated in memory and learning processes. Over 40% of neuronal synapses are glutamatergic. The glutamatergic system is thought to play a fundamental role in long-term potentiation (LTP), a mechanism underlying learning and memory storage in the hippocampus and neocortex. Deficient levels of glutamate have been reported in brains of AD patients [40]. Loss of glutamate activity correlates to clinical dementia and MCI [41], playing a role in the pathogenesis of AD since early states [42]. This decrease in glutamate especially impacts on the posterior cingulate cortex, a region involved in cerebral glucose metabolism and beta-amyloid deposition in MCI [41]. The data suggest that the musical stimuli could contribute to managing L-glutamate in the brains of ACD patients, a target shared with traditional pharmacological therapies. Thus, among the most common current treatments aimed at reducing cognitive decline in AD are those targeting the cholinesterase and glutamatergic systems [43].

*Sphingolipidome and regulation of the Aβ peptides*

Another important difference between ACD and healthy controls is in the metabolism of sphingolipids. The complex universe of the sphingolipidome regulates a plethora of biological processes in cells. These molecules have kinases, phosphatase, and lipases as main targets to exert distinct cellular functions, and they have been found to be associated with neurodegenerative processes, as well as metabolic disorders, cancer, immune functions, and cardiovascular disorders [44]. An adequate regulation of sphingolipids metabolism in the brain is essential, such that subtle deviations of the sphingolipids balance might be sufficient to produce neurodegenerative conditions, such as AD or Parkinson [45]. Sphingolipids alteration can contribute to the neuropathological features of AD, including Aβ production, Tau formation and neurodegeneration. In this regard, sphingolipids metabolism correlated with Aβ levels in cerebrospinal fluid from AD patients [46] strengthening this effect of the sphingolipids balance disturbance in the Aβ clearance. Aβ in AD can produce membrane oxidative stress, resulting in accumulation of long-chain ceramides and cholesterol unleashing the clinical manifestations of the disease [47].

The accumulation of Aβ in the brain is an early toxic event in the pathogenesis of AD, and there is no efficient pharmacological treatment to stop it. The amyloid cascade hypothesis states that Aβ accumulation triggers tau hyperphosphorylation and aggregation; ultimately, these aggregates lead to inflammation, synaptic impairment, neuronal loss, and thus cognitive decline and behavioral abnormalities [48]. Even though our methodological approach cannot shed light on why music stimuli target on the regulation of Aβ (increasing the expression in ACD patients >15 times the expression of controls), it is a striking observation that music is able to impact on this key signature of neurodegeneration in AD patients. It is mandatory to investigate the relationship between this observation and the benefits of music reported by others in AD patients [9].

Co-expression networks

*Autophagic impairment*

Autophagic impairment has been related to neurodegenerative processes since it regulates intracytoplasmic levels of aggregate-prone proteins responsible for common neurodegenerative diseases, such as AD, Parkinson disease, Huntington disease, amyotrophic lateral sclerosis, or dementia [49-52]. Aggregates of Tau and Aβ as well as accumulation of autophagosomes in neurons of AD patients are clear signals of autophagic system disruption. Autophagy is a key regulator of Aβ generation and clearance, as Aβ is released from neurons through an autophagy-dependent mechanism [53]. Moreover, accumulation of autophagosomes (autophagosomes and autolysosomes [autophagic vacuoles]) has been observed in neurons from AD patients in different stages [54, 55]. However, there are discrepancies in the literature as to whether an up-regulation of autophagic activity is responsible for the accumulation of these autophagosomes [56] or, conversely, whether this accumulation might be due to a decrease in autophagic-mediated protein release or autophagosome clearance activity [55, 57, 58]. Recent studies claim that autophagy can be differentially affected at different stages of AD [53, 59]. Currently, different autophagy modulators are being investigated as treatments for AD [60, 61].

Enrichment map

The Receptor for advanced glycation end products (RAGE) is a multiligand transmembrane receptor of the immunoglobulin superfamily. It is highly expressed in neurons of patients suffering neurodegenerative diseases and is also significantly up-regulated in AD-related optic neuropathy [62]. RAGE transports the Aβ toxins through the blood-brain barrier; however, in AD, RAGE mediates Aβ toxicity, disrupting the blood-brain barrier and promoting AD occurrence and progression [63, 64]. The up-regulation of RAGE accelerates Aβ uptake and transport, facilitating blood-brain barrier crossing *via* endocytosis, activating inflammatory pathways, oxidative stress, neuroinflammation, and synaptic and neuronal destruction [65, 66]. Moreover, accumulation of Aβ oligomers can also induce RAGE-mediated autophagy, interfering with the proper functioning of tight junction proteins in the blood-brain barrier and, thus, contributing to the worsening of AD [67, 68]. Due to its role in MCI/AD pathogenesis, RAGE is being investigated as a therapeutic target in AD [69]; reviewed in [70].

Limitations

There are various limitations to the present study. First, its design does not allow to completely isolate the music stimuli from other possible stimuli and confounding factors. This fact, however, characterizes most (if not all) studies targeting complex multifactorial traits e.g. [71-74]. It is the population-based nature of these discovery-based studies that minimizes the possible impact of individual features in transcriptomes (which could deviate from the ‘average’ due to possible confounded factors). Moreover, to minimize possible confounding effects, the experimental concert Sensogenomics22-pilot was conceived to reduce potential noise factors as much as possible. For instance, the procedures for sample collection were minimally invasive (capillary puncture), with the aim of reducing possible stress factors, e.g., avoiding waiting times for patients that could develop anxiety or stress, accompaniment of patients by their caregivers, etc. In this regard, it has been reported that transcriptomes analyzed from fingerstick (capillary) blood samples (as done here) represent a very good proxy for transcriptomes analyzed from venipuncture (venous) blood [75, 76], although this issue might be controversial, at least in the context of vaccination [77]. In any case, it seems clear from our results that capillary blood samples capture transcriptomics changes occurring because of the simple impact of a sensorial stimulus. These changes are most likely echoing those occurring in the brain, as this is the natural anatomical entry point for the musical stimuli. In addition, the musical event took place in a natural place for music, as it would with any other non-experimental musical event. The ecological validity of the experiment adds extra value to the study; to the best of our knowledge, there is no comparable precedent in the scientific literature. Among other limitations, it is noteworthy to mention that, although we have investigated many more samples than previous experiments on the effect of music on transcriptomes, it is mandatory to validate them using much larger cohorts. Finally, we have detected more changes in the transcriptomes of ACD patients than in the transcriptomes of controls; given that the sample size of both cohorts was comparable, it is highly probable that the musical stimuli might have a higher impact on the transcriptomes of cases than in those from controls. A powerful study would be needed to validate the results of the present study and increase the probability to detect more changes occurring in the transcriptomes of healthy controls.

**Supplementary Text Figure 1.** Soft-thresholding power estimation in ACD and healthy controls cohorts. Plots represent correlation between soft-thresholding powers and both the scale-free fit index and the mean connectivity.

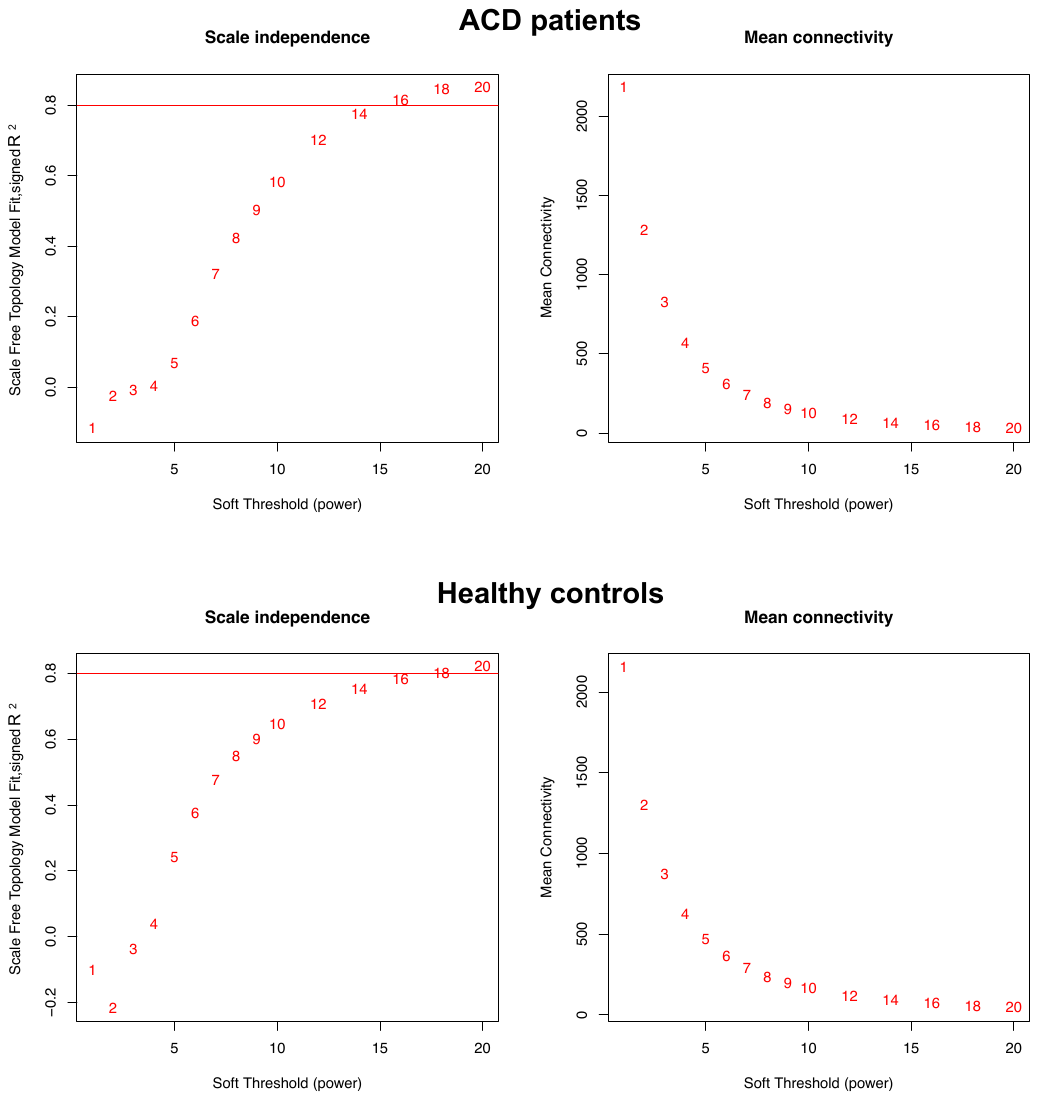

**Supplementary Text Figure 2**. (A) Music-related DEGs in ACD patients when comparing TP1 *vs*. TP2 (adjusted *P*-value <0.05), and (B) DEGs involved in neuronal-related processes (*P*-values < 10^-5^) in ACD patients and controls when comparing when comparing TP1 *vs*. TP2.

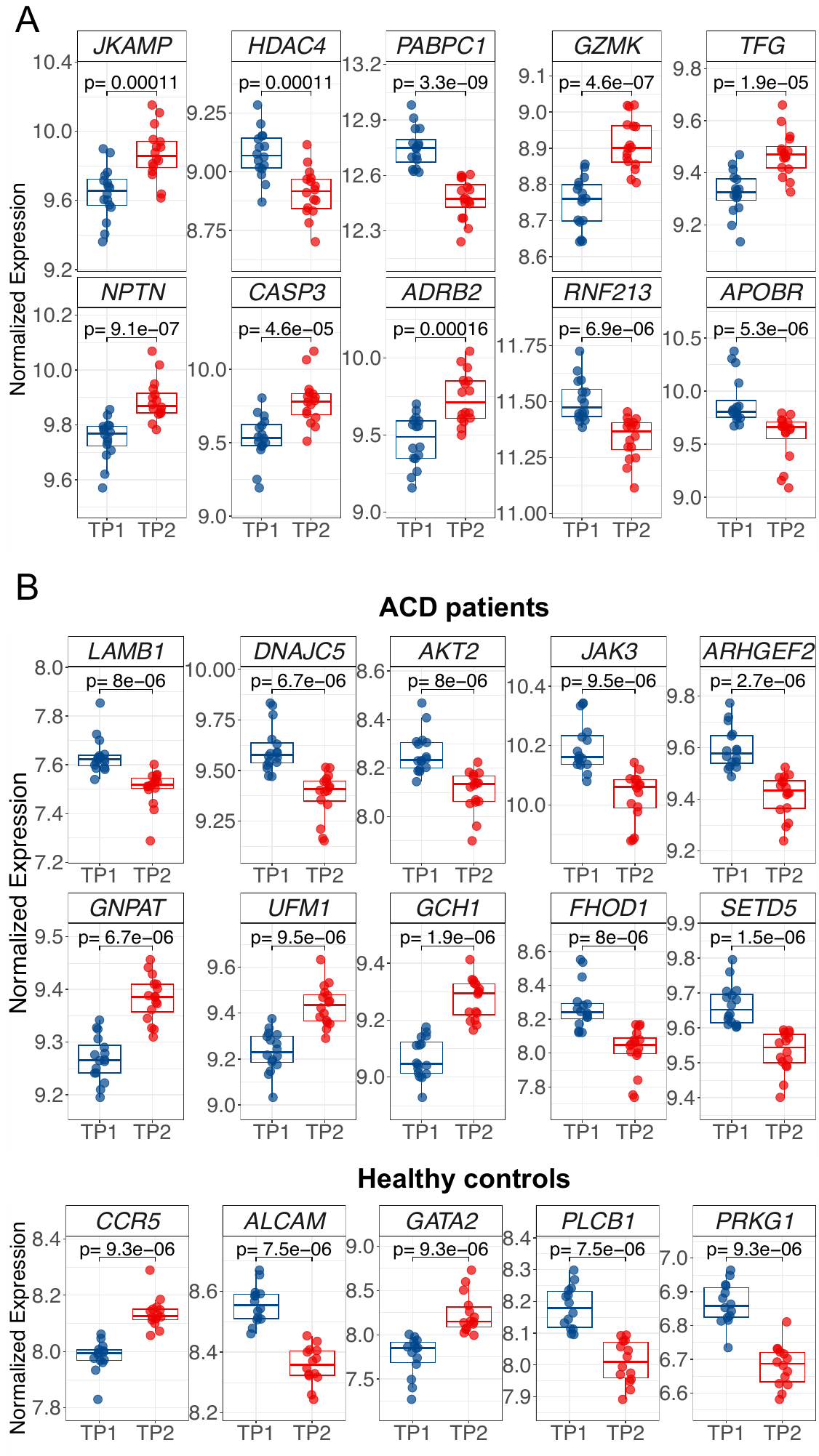

**Supplementary Text Figure 3.** Co-expression network analysis in healthy control cohort. (A) Clustering dendrogram of genes and co-expression modules detected represented by different colors. (B) Correlation values heatmap obtained from WGCNA module analysis of co-expressed genes in healthy controls. Upper value shows to individual correlation value of the module with musical stimuli. *P*-values of these correlations are represented in brackets (lower values). (C) Hierarchical clustering eigengene dendrogram and heatmap for healthy controls dataset showing relationships among the modules and TP (musical stimuli; TP2 *vs.* TP1). Gene names on the left of the heatmap are the hub genes of each module. (D) Raincloud plots of differences in samples eigengene values between TP1 and TP2 from modules showing the largest statistical significance in healthy controls. (E) Plot showing comparison between MM (module membership) and musical stimuli correlation of genes from the most significant module detected in the healthy control cohort.

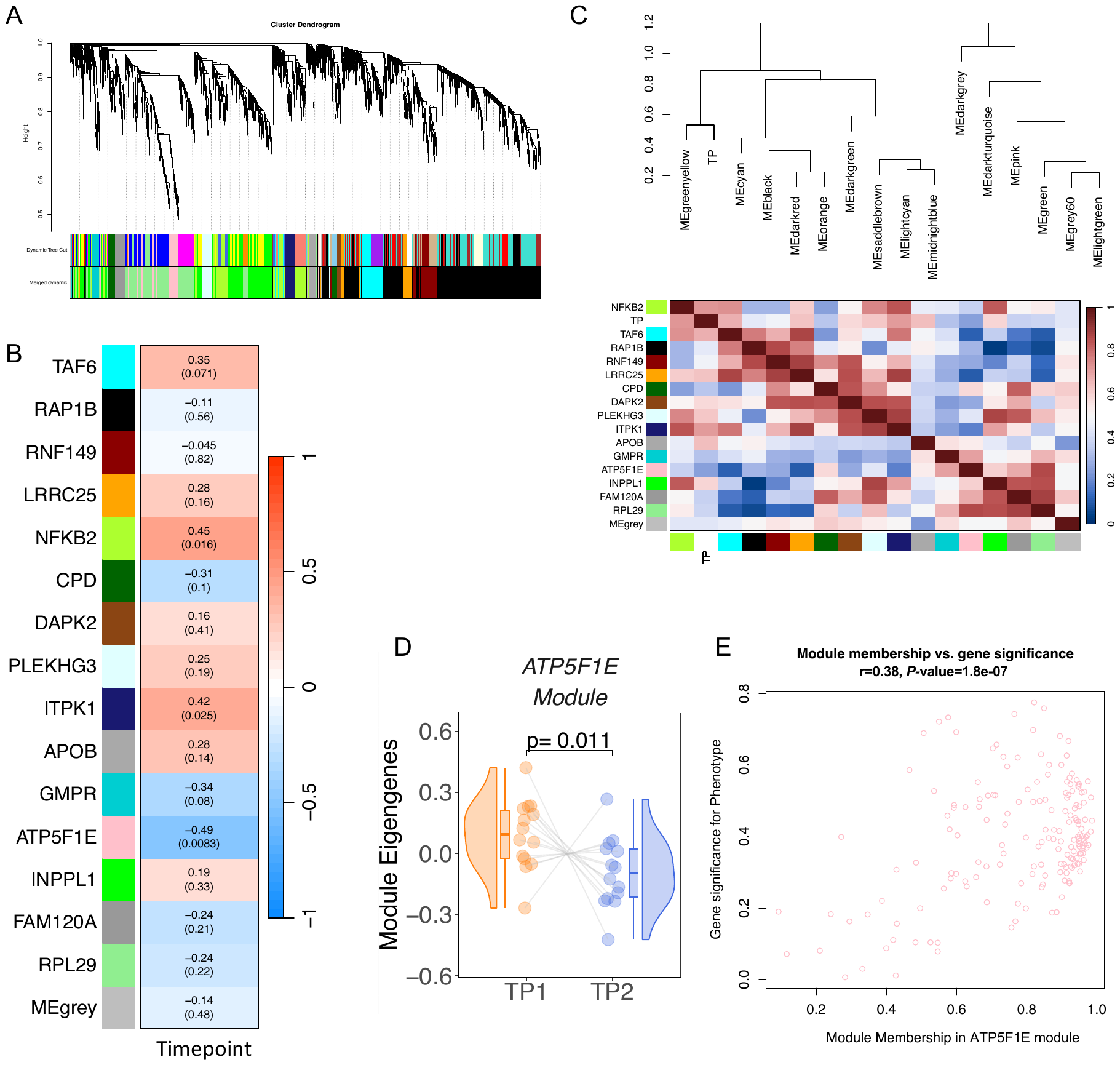

**Supplementary Text Figure 4.** (A) Top GO biological processes detected for the *ATP5F1E* and *ITPK1* modules from healthy controls. (B) Module membership (MM) and phenotype correlation (musical stimuli) for genes from the most significant pathways obtained from the module *ATP5F1E*.

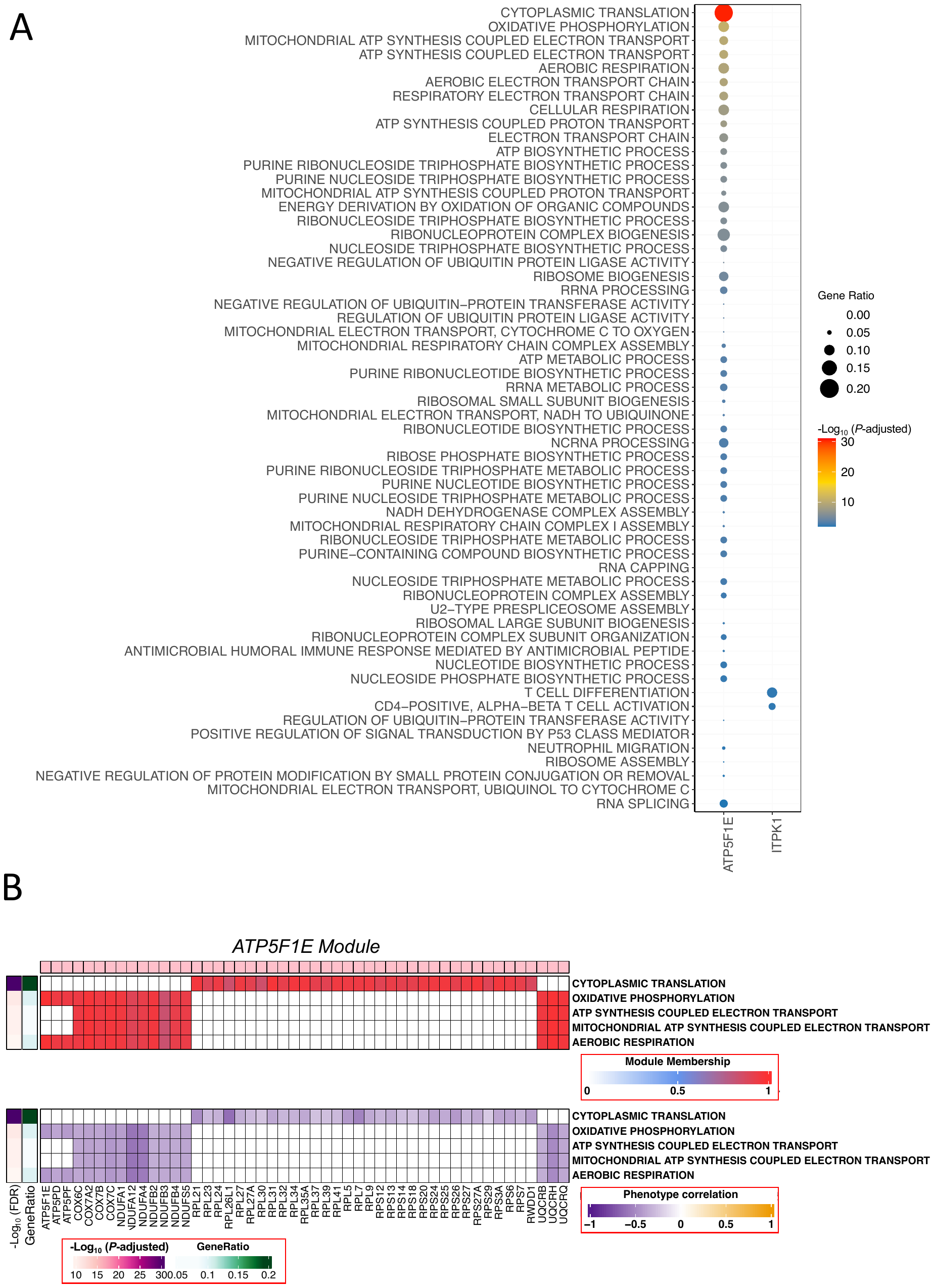

**Supplementary Text Table 1.** Neuro-biological related terms retrieved from Gene Ontology database (GO).

**GO terms**

ADENYLATE CYCLASE ACTIVATING ADRENERGIC RECEPTOR SIGNALING PATHWAY

ADENYLATE CYCLASE ACTIVATING DOPAMINE RECEPTOR SIGNALING PATHWAY

ADRENAL GLAND DEVELOPMENT

ADRENERGIC RECEPTOR SIGNALING PATHWAY

AMINO ACID NEUROTRANSMITTER REUPTAKE

AMPA GLUTAMATE RECEPTOR CLUSTERING

AMYLOID BETA CLEARANCE

AMYLOID BETA METABOLIC PROCESS

AMYLOID FIBRIL FORMATION

AMYLOID PRECURSOR PROTEIN BIOSYNTHETIC PROCESS

AMYLOID PRECURSOR PROTEIN CATABOLIC PROCESS

AMYLOID PRECURSOR PROTEIN METABOLIC PROCESS

ASTROCYTE ACTIVATION

ASTROCYTE DEVELOPMENT

ASTROCYTE DIFFERENTIATION

AUTONOMIC NERVOUS SYSTEM DEVELOPMENT

AXO DENDRITIC TRANSPORT

AXON ENSHEATHMENT IN CENTRAL NERVOUS SYSTEM

BERGMANN GLIAL CELL DIFFERENTIATION

BRAIN MORPHOGENESIS

BRANCHED CHAIN AMINO ACID CATABOLIC PROCESS

BRANCHED CHAIN AMINO ACID METABOLIC PROCESS

BRANCHED CHAIN AMINO ACID TRANSPORT

BRANCHING MORPHOGENESIS OF A NERVE

BUNDLE OF HIS CELL TO PURKINJE MYOCYTE COMMUNICATION

CALCIUM ION REGULATED EXOCYTOSIS OF NEUROTRANSMITTER

CARDIAC NEURAL CREST CELL DEVELOPMENT INVOLVED IN OUTFLOW TRACT MORPHOGENESIS

CARDIAC NEURAL CREST CELL DIFFERENTIATION INVOLVED IN HEART DEVELOPMENT

CARDIAC NEURAL CREST CELL MIGRATION INVOLVED IN OUTFLOW TRACT MORPHOGENESIS

CATECHOLAMINE UPTAKE INVOLVED IN SYNAPTIC TRANSMISSION

CELL DIFFERENTIATION IN HINDBRAIN

CELL MIGRATION IN HINDBRAIN

CELL MORPHOGENESIS INVOLVED IN NEURON DIFFERENTIATION

CELL PROLIFERATION IN FOREBRAIN

CELL PROLIFERATION IN HINDBRAIN

CELLULAR RESPONSE TO AMYLOID BETA

CELLULAR RESPONSE TO CORTICOSTEROID STIMULUS

CELLULAR RESPONSE TO EPINEPHRINE STIMULUS

CELLULAR RESPONSE TO LEPTIN STIMULUS

CELLULAR RESPONSE TO MINERALOCORTICOID STIMULUS

CENTRAL NERVOUS SYSTEM DEVELOPMENT

CENTRAL NERVOUS SYSTEM NEURON AXONOGENESIS

CENTRAL NERVOUS SYSTEM NEURON DEVELOPMENT

CENTRAL NERVOUS SYSTEM NEURON DIFFERENTIATION

CENTRAL NERVOUS SYSTEM PROJECTION NEURON AXONOGENESIS

CERAMIDE BIOSYNTHETIC PROCESS

CERAMIDE CATABOLIC PROCESS

CERAMIDE METABOLIC PROCESS

CERAMIDE TRANSPORT

CEREBELLAR CORTEX DEVELOPMENT

CEREBELLAR CORTEX FORMATION

CEREBELLAR CORTEX MORPHOGENESIS

CEREBELLAR GRANULAR LAYER DEVELOPMENT

CEREBELLAR PURKINJE CELL LAYER DEVELOPMENT

CEREBELLAR PURKINJE CELL LAYER FORMATION

CEREBELLAR PURKINJE CELL LAYER MORPHOGENESIS

CEREBRAL CORTEX CELL MIGRATION

CEREBRAL CORTEX DEVELOPMENT

CEREBRAL CORTEX GABAERGIC INTERNEURON DIFFERENTIATION

CEREBRAL CORTEX NEURON DIFFERENTIATION

CEREBRAL CORTEX RADIALLY ORIENTED CELL MIGRATION

CEREBROSPINAL FLUID CIRCULATION

CIRCADIAN SLEEP WAKE CYCLE

CIRCADIAN SLEEP WAKE CYCLE SLEEP

COGNITION

COMMISSURAL NEURON AXON GUIDANCE

CORTICAL ACTIN CYTOSKELETON ORGANIZATION

CORTICAL CYTOSKELETON ORGANIZATION

CORTICOSTEROID HORMONE SECRETION

CORTICOSTEROID RECEPTOR SIGNALING PATHWAY

CRANIAL NERVE DEVELOPMENT

CRANIAL NERVE FORMATION

CRANIAL NERVE MORPHOGENESIS

CRANIAL NERVE STRUCTURAL ORGANIZATION

DENDRITE DEVELOPMENT

DENDRITE EXTENSION

DENDRITE MORPHOGENESIS

DENDRITE SELF AVOIDANCE

DENDRITIC CELL ANTIGEN PROCESSING AND PRESENTATION

DENDRITIC CELL APOPTOTIC PROCESS

DENDRITIC CELL CHEMOTAXIS

DENDRITIC CELL CYTOKINE PRODUCTION

DENDRITIC CELL DIFFERENTIATION

DENDRITIC CELL MIGRATION

DENDRITIC SPINE DEVELOPMENT

DENDRITIC SPINE MAINTENANCE

DENDRITIC SPINE MORPHOGENESIS

DENDRITIC TRANSPORT

DOPAMINE BIOSYNTHETIC PROCESS

DOPAMINE METABOLIC PROCESS

DOPAMINE RECEPTOR SIGNALING PATHWAY

DOPAMINE SECRETION

DOPAMINE TRANSPORT

DOPAMINERGIC NEURON DIFFERENTIATION

DORSAL VENTRAL NEURAL TUBE PATTERNING

EMBRYONIC BRAIN DEVELOPMENT

EMBRYONIC FORELIMB MORPHOGENESIS

EMBRYONIC HINDLIMB MORPHOGENESIS

ENSHEATHMENT OF NEURONS

ENTERIC NERVOUS SYSTEM DEVELOPMENT

EPINEPHRINE TRANSPORT

ESTABLISHMENT OF BLOOD BRAIN BARRIER

ESTABLISHMENT OF PLANAR POLARITY INVOLVED IN NEURAL TUBE CLOSURE

EXCITATORY SYNAPSE ASSEMBLY

FACIAL NERVE MORPHOGENESIS

FOREBRAIN CELL MIGRATION

FOREBRAIN DEVELOPMENT

FOREBRAIN GENERATION OF NEURONS

FOREBRAIN MORPHOGENESIS

FOREBRAIN NEURON DEVELOPMENT

FOREBRAIN NEURON DIFFERENTIATION

FOREBRAIN REGIONALIZATION

FORELIMB MORPHOGENESIS

G PROTEIN COUPLED ACETYLCHOLINE RECEPTOR SIGNALING PATHWAY

G PROTEIN COUPLED GLUTAMATE RECEPTOR SIGNALING PATHWAY

GABAERGIC NEURON DIFFERENTIATION

GAMMA AMINOBUTYRIC ACID SECRETION

GAMMA AMINOBUTYRIC ACID SIGNALING PATHWAY

GAMMA AMINOBUTYRIC ACID TRANSPORT

GANGLIOSIDE BIOSYNTHETIC PROCESS

GANGLIOSIDE METABOLIC PROCESS

GENERATION OF NEURONS

GLIAL CELL ACTIVATION

GLIAL CELL APOPTOTIC PROCESS

GLIAL CELL DERIVED NEUROTROPHIC FACTOR RECEPTOR SIGNALING PATHWAY

GLIAL CELL DEVELOPMENT

GLIAL CELL DIFFERENTIATION

GLIAL CELL FATE COMMITMENT

GLIAL CELL MIGRATION

GLIAL CELL PROLIFERATION

GLUCOCORTICOID BIOSYNTHETIC PROCESS

GLUCOCORTICOID METABOLIC PROCESS

GLUCOCORTICOID SECRETION

GLUTAMATE METABOLIC PROCESS

GLUTAMATE RECEPTOR SIGNALING PATHWAY

GLUTAMATE SECRETION

GLUTAMINE FAMILY AMINO ACID BIOSYNTHETIC PROCESS

GLUTAMINE FAMILY AMINO ACID CATABOLIC PROCESS

GLUTAMINE FAMILY AMINO ACID METABOLIC PROCESS

GLUTAMINE METABOLIC PROCESS

GLYCOSPHINGOLIPID BIOSYNTHETIC PROCESS

GLYCOSPHINGOLIPID METABOLIC PROCESS

GLYCOSYLCERAMIDE METABOLIC PROCESS

HINDBRAIN DEVELOPMENT

HINDBRAIN MORPHOGENESIS

HINDLIMB MORPHOGENESIS

HIPPOCAMPUS DEVELOPMENT

HISTAMINE PRODUCTION INVOLVED IN INFLAMMATORY RESPONSE

HISTAMINE SECRETION

HISTAMINE TRANSPORT

HYPOTHALAMUS CELL DIFFERENTIATION

HYPOTHALAMUS DEVELOPMENT

IMMUNOLOGICAL MEMORY FORMATION PROCESS

IMMUNOLOGICAL MEMORY PROCESS

IMMUNOLOGICAL SYNAPSE FORMATION

INHIBITORY POSTSYNAPTIC POTENTIAL

INHIBITORY SYNAPSE ASSEMBLY

INNERVATION

INTERNEURON MIGRATION

L GLUTAMATE IMPORT ACROSS PLASMA MEMBRANE

L GLUTAMATE TRANSMEMBRANE TRANSPORT

LAYER FORMATION IN CEREBRAL CORTEX

LEPTIN MEDIATED SIGNALING PATHWAY

LIMB BUD FORMATION

LIMBIC SYSTEM DEVELOPMENT

LONG TERM MEMORY

LONG TERM SYNAPTIC DEPRESSION

LONG TERM SYNAPTIC POTENTIATION

MAINTENANCE OF BLOOD BRAIN BARRIER

MAINTENANCE OF SYNAPSE STRUCTURE

MEMORY

MIDBRAIN DEVELOPMENT

MIDBRAIN DOPAMINERGIC NEURON DIFFERENTIATION

MINERALOCORTICOID BIOSYNTHETIC PROCESS

MINERALOCORTICOID METABOLIC PROCESS

MODIFICATION OF POSTSYNAPTIC STRUCTURE

MODIFICATION OF SYNAPTIC STRUCTURE

MODULATION OF EXCITATORY POSTSYNAPTIC POTENTIAL

MOTOR NEURON APOPTOTIC PROCESS

MOTOR NEURON AXON GUIDANCE

MOTOR NEURON MIGRATION

MYELIN ASSEMBLY

MYELIN MAINTENANCE

MYELOID DENDRITIC CELL ACTIVATION

MYELOID DENDRITIC CELL DIFFERENTIATION

NEGATIVE REGULATION OF AMYLOID FIBRIL FORMATION

NEGATIVE REGULATION OF AMYLOID PRECURSOR PROTEIN CATABOLIC PROCESS

NEGATIVE REGULATION OF ASTROCYTE DIFFERENTIATION

NEGATIVE REGULATION OF DENDRITIC SPINE DEVELOPMENT

NEGATIVE REGULATION OF GLIAL CELL DIFFERENTIATION

NEGATIVE REGULATION OF GLIAL CELL PROLIFERATION

NEGATIVE REGULATION OF LONG TERM SYNAPTIC POTENTIATION

NEGATIVE REGULATION OF MOTOR NEURON APOPTOTIC PROCESS

NEGATIVE REGULATION OF MYELINATION

NEGATIVE REGULATION OF NERVOUS SYSTEM DEVELOPMENT

NEGATIVE REGULATION OF NERVOUS SYSTEM PROCESS

NEGATIVE REGULATION OF NEURAL PRECURSOR CELL PROLIFERATION

NEGATIVE REGULATION OF NEUROINFLAMMATORY RESPONSE

NEGATIVE REGULATION OF NEURON APOPTOTIC PROCESS

NEGATIVE REGULATION OF NEURON DEATH

NEGATIVE REGULATION OF NEURON DIFFERENTIATION

NEGATIVE REGULATION OF NEURON MIGRATION

NEGATIVE REGULATION OF NEURON PROJECTION DEVELOPMENT

NEGATIVE REGULATION OF NEURON PROJECTION REGENERATION

NEGATIVE REGULATION OF NEUROTRANSMITTER SECRETION

NEGATIVE REGULATION OF NEUROTRANSMITTER TRANSPORT

NEGATIVE REGULATION OF OLIGODENDROCYTE DIFFERENTIATION

NEGATIVE REGULATION OF OXIDATIVE STRESS INDUCED NEURON DEATH

NEGATIVE REGULATION OF SYNAPSE ORGANIZATION

NEGATIVE REGULATION OF SYNAPTIC TRANSMISSION

NEGATIVE REGULATION OF SYNAPTIC TRANSMISSION GLUTAMATERGIC

NERVE DEVELOPMENT

NERVE GROWTH FACTOR SIGNALING PATHWAY

NERVOUS SYSTEM PROCESS

NERVOUS SYSTEM PROCESS INVOLVED IN REGULATION OF SYSTEMIC ARTERIAL BLOOD PRESSURE

NEURAL CREST CELL DIFFERENTIATION

NEURAL CREST FORMATION

NEURAL NUCLEUS DEVELOPMENT

NEURAL PRECURSOR CELL PROLIFERATION

NEURAL RETINA DEVELOPMENT

NEURAL TUBE DEVELOPMENT

NEURAL TUBE FORMATION

NEURAL TUBE PATTERNING

NEUROBLAST DIVISION

NEUROBLAST PROLIFERATION

NEUROENDOCRINE CELL DIFFERENTIATION

NEUROEPITHELIAL CELL DIFFERENTIATION

NEUROGENESIS

NEUROINFLAMMATORY RESPONSE

NEUROMUSCULAR JUNCTION DEVELOPMENT

NEUROMUSCULAR PROCESS

NEUROMUSCULAR PROCESS CONTROLLING BALANCE

NEUROMUSCULAR PROCESS CONTROLLING POSTURE

NEUROMUSCULAR SYNAPTIC TRANSMISSION

NEURON APOPTOTIC PROCESS

NEURON CELL CELL ADHESION

NEURON CELLULAR HOMEOSTASIS

NEURON DEATH

NEURON DEATH IN RESPONSE TO OXIDATIVE STRESS

NEURON DEVELOPMENT

NEURON FATE COMMITMENT

NEURON FATE DETERMINATION

NEURON FATE SPECIFICATION

NEURON MATURATION

NEURON MIGRATION

NEURON PROJECTION ARBORIZATION

NEURON PROJECTION EXTENSION

NEURON PROJECTION EXTENSION INVOLVED IN NEURON PROJECTION GUIDANCE

NEURON PROJECTION GUIDANCE

NEURON PROJECTION MAINTENANCE

NEURON PROJECTION ORGANIZATION

NEURON PROJECTION REGENERATION

NEURON RECOGNITION

NEURON REMODELING

NEURONAL ACTION POTENTIAL

NEURONAL ION CHANNEL CLUSTERING

NEURONAL STEM CELL POPULATION MAINTENANCE

NEUROPEPTIDE SIGNALING PATHWAY

NEUROTRANSMITTER BIOSYNTHETIC PROCESS

NEUROTRANSMITTER CATABOLIC PROCESS

NEUROTRANSMITTER GATED ION CHANNEL CLUSTERING

NEUROTRANSMITTER METABOLIC PROCESS

NEUROTRANSMITTER RECEPTOR INTERNALIZATION

NEUROTRANSMITTER RECEPTOR LOCALIZATION TO POSTSYNAPTIC SPECIALIZATION MEMBRANE

NEUROTRANSMITTER RECEPTOR TRANSPORT

NEUROTRANSMITTER RECEPTOR TRANSPORT ENDOSOME TO POSTSYNAPTIC MEMBRANE

NEUROTRANSMITTER RECEPTOR TRANSPORT TO PLASMA MEMBRANE

NEUROTRANSMITTER REUPTAKE

NEUROTRANSMITTER SECRETION

NEUROTRANSMITTER TRANSPORT

NEUROTRANSMITTER UPTAKE

NEUROTROPHIN SIGNALING PATHWAY

NEUROTROPHIN TRK RECEPTOR SIGNALING PATHWAY

NORADRENERGIC NEURON DIFFERENTIATION

NOREPINEPHRINE METABOLIC PROCESS

NOREPINEPHRINE SECRETION

NOREPINEPHRINE TRANSPORT

OLFACTORY BULB INTERNEURON DIFFERENTIATION

OLIGODENDROCYTE DEVELOPMENT

OLIGODENDROCYTE DIFFERENTIATION

OPTIC NERVE DEVELOPMENT

PARASYMPATHETIC NERVOUS SYSTEM DEVELOPMENT

PEPTIDYL GLUTAMIC ACID MODIFICATION

PERIPHERAL NERVOUS SYSTEM DEVELOPMENT

PERIPHERAL NERVOUS SYSTEM NEURON DIFFERENTIATION

PITUITARY GLAND DEVELOPMENT

PLANAR CELL POLARITY PATHWAY INVOLVED IN NEURAL TUBE CLOSURE

POSITIVE REGULATION OF AMYLOID BETA FORMATION

POSITIVE REGULATION OF AMYLOID PRECURSOR PROTEIN CATABOLIC PROCESS

POSITIVE REGULATION OF ASTROCYTE DIFFERENTIATION

POSITIVE REGULATION OF DENDRITE DEVELOPMENT

POSITIVE REGULATION OF DENDRITE MORPHOGENESIS

POSITIVE REGULATION OF DENDRITIC CELL CHEMOTAXIS

POSITIVE REGULATION OF DENDRITIC CELL CYTOKINE PRODUCTION

POSITIVE REGULATION OF DENDRITIC SPINE DEVELOPMENT

POSITIVE REGULATION OF DENDRITIC SPINE MORPHOGENESIS

POSITIVE REGULATION OF EXCITATORY POSTSYNAPTIC POTENTIAL

POSITIVE REGULATION OF GLIAL CELL DIFFERENTIATION

POSITIVE REGULATION OF GLIAL CELL MIGRATION

POSITIVE REGULATION OF GLIAL CELL PROLIFERATION

POSITIVE REGULATION OF GLUTAMATE SECRETION

POSITIVE REGULATION OF LONG TERM SYNAPTIC POTENTIATION

POSITIVE REGULATION OF MYELINATION

POSITIVE REGULATION OF NERVOUS SYSTEM DEVELOPMENT

POSITIVE REGULATION OF NERVOUS SYSTEM PROCESS

POSITIVE REGULATION OF NEURAL PRECURSOR CELL PROLIFERATION

POSITIVE REGULATION OF NEUROBLAST PROLIFERATION

POSITIVE REGULATION OF NEUROGENESIS

POSITIVE REGULATION OF NEUROINFLAMMATORY RESPONSE

POSITIVE REGULATION OF NEURON APOPTOTIC PROCESS

POSITIVE REGULATION OF NEURON DEATH

POSITIVE REGULATION OF NEURON DIFFERENTIATION

POSITIVE REGULATION OF NEURON MIGRATION

POSITIVE REGULATION OF NEURON PROJECTION DEVELOPMENT

POSITIVE REGULATION OF NEURON PROJECTION REGENERATION

POSITIVE REGULATION OF NEUROTRANSMITTER SECRETION

POSITIVE REGULATION OF NEUROTRANSMITTER TRANSPORT

POSITIVE REGULATION OF OLIGODENDROCYTE DIFFERENTIATION

POSITIVE REGULATION OF SYNAPSE ASSEMBLY

POSITIVE REGULATION OF SYNAPTIC TRANSMISSION

POSITIVE REGULATION OF SYNAPTIC TRANSMISSION GABAERGIC

POSITIVE REGULATION OF SYNAPTIC TRANSMISSION GLUTAMATERGIC

POSTSYNAPSE ASSEMBLY

POSTSYNAPSE ORGANIZATION

POSTSYNAPTIC ACTIN CYTOSKELETON ORGANIZATION

POSTSYNAPTIC CYTOSKELETON ORGANIZATION

POSTSYNAPTIC MEMBRANE ASSEMBLY

POSTSYNAPTIC MEMBRANE ORGANIZATION

POSTSYNAPTIC MODULATION OF CHEMICAL SYNAPTIC TRANSMISSION

POSTSYNAPTIC NEUROTRANSMITTER RECEPTOR INTERNALIZATION

POSTSYNAPTIC SIGNAL TRANSDUCTION

POSTSYNAPTIC SPECIALIZATION ASSEMBLY

POSTSYNAPTIC SPECIALIZATION ORGANIZATION

PRESYNAPSE ORGANIZATION

PRESYNAPTIC ENDOCYTOSIS

PRESYNAPTIC MODULATION OF CHEMICAL SYNAPTIC TRANSMISSION

PROTEIN LOCALIZATION TO CELL CORTEX

PROTEIN LOCALIZATION TO POSTSYNAPSE

PROTEIN LOCALIZATION TO PRESYNAPSE

PROTEIN LOCALIZATION TO SYNAPSE

PROTEIN POLYGLUTAMYLATION

PYRAMIDAL NEURON DEVELOPMENT

PYRAMIDAL NEURON DIFFERENTIATION

RADIAL GLIAL CELL DIFFERENTIATION

RECEPTOR LOCALIZATION TO SYNAPSE

REGULATION OF AMYLOID BETA CLEARANCE

REGULATION OF AMYLOID FIBRIL FORMATION

REGULATION OF AMYLOID PRECURSOR PROTEIN CATABOLIC PROCESS

REGULATION OF ASPARTIC TYPE ENDOPEPTIDASE ACTIVITY INVOLVED IN AMYLOID PRECURSOR PROTEIN CATABOLIC PROCESS

REGULATION OF ASTROCYTE DIFFERENTIATION

REGULATION OF CEREBELLAR GRANULE CELL PRECURSOR PROLIFERATION

REGULATION OF CIRCADIAN SLEEP WAKE CYCLE

REGULATION OF CORTICOSTEROID HORMONE SECRETION

REGULATION OF DENDRITE DEVELOPMENT

REGULATION OF DENDRITE EXTENSION

REGULATION OF DENDRITE MORPHOGENESIS

REGULATION OF DENDRITIC CELL ANTIGEN PROCESSING AND PRESENTATION

REGULATION OF DENDRITIC CELL CHEMOTAXIS

REGULATION OF DENDRITIC CELL DIFFERENTIATION

REGULATION OF DENDRITIC SPINE DEVELOPMENT

REGULATION OF DENDRITIC SPINE MAINTENANCE

REGULATION OF DENDRITIC SPINE MORPHOGENESIS

REGULATION OF DOPAMINE RECEPTOR SIGNALING PATHWAY

REGULATION OF DOPAMINERGIC NEURON DIFFERENTIATION

REGULATION OF EXCITATORY SYNAPSE ASSEMBLY

REGULATION OF GLIAL CELL APOPTOTIC PROCESS

REGULATION OF GLIAL CELL DIFFERENTIATION

REGULATION OF GLIAL CELL MIGRATION

REGULATION OF GLIAL CELL PROLIFERATION

REGULATION OF GLUCOCORTICOID METABOLIC PROCESS

REGULATION OF GLUCOCORTICOID SECRETION

REGULATION OF GLUTAMATE SECRETION

REGULATION OF LONG TERM NEURONAL SYNAPTIC PLASTICITY

REGULATION OF LONG TERM SYNAPTIC DEPRESSION

REGULATION OF LONG TERM SYNAPTIC POTENTIATION

REGULATION OF MICROGLIAL CELL ACTIVATION

REGULATION OF MOTOR NEURON APOPTOTIC PROCESS

REGULATION OF MYELINATION

REGULATION OF NERVOUS SYSTEM DEVELOPMENT

REGULATION OF NERVOUS SYSTEM PROCESS

REGULATION OF NEURAL PRECURSOR CELL PROLIFERATION

REGULATION OF NEUROBLAST PROLIFERATION

REGULATION OF NEUROGENESIS

REGULATION OF NEUROINFLAMMATORY RESPONSE

REGULATION OF NEURON DIFFERENTIATION

REGULATION OF NEURON MIGRATION

REGULATION OF NEURON PROJECTION ARBORIZATION

REGULATION OF NEURON PROJECTION DEVELOPMENT

REGULATION OF NEURON PROJECTION REGENERATION

REGULATION OF NEURONAL SYNAPTIC PLASTICITY

REGULATION OF NEUROTRANSMITTER LEVELS

REGULATION OF NEUROTRANSMITTER RECEPTOR ACTIVITY

REGULATION OF NEUROTRANSMITTER TRANSPORT

REGULATION OF NEUROTRANSMITTER UPTAKE

REGULATION OF NEUROTROPHIN TRK RECEPTOR SIGNALING PATHWAY

REGULATION OF OLIGODENDROCYTE DIFFERENTIATION

REGULATION OF POSTSYNAPSE ORGANIZATION

REGULATION OF POSTSYNAPTIC CYTOSOLIC CALCIUM ION CONCENTRATION

REGULATION OF POSTSYNAPTIC DENSITY ASSEMBLY

REGULATION OF POSTSYNAPTIC DENSITY ORGANIZATION

REGULATION OF POSTSYNAPTIC MEMBRANE NEUROTRANSMITTER RECEPTOR LEVELS

REGULATION OF POSTSYNAPTIC MEMBRANE POTENTIAL

REGULATION OF POSTSYNAPTIC NEUROTRANSMITTER RECEPTOR ACTIVITY

REGULATION OF POSTSYNAPTIC NEUROTRANSMITTER RECEPTOR INTERNALIZATION

REGULATION OF POSTSYNAPTIC SPECIALIZATION ASSEMBLY

REGULATION OF PRESYNAPSE ORGANIZATION

REGULATION OF PRESYNAPTIC CYTOSOLIC CALCIUM ION CONCENTRATION

REGULATION OF PROTEIN LOCALIZATION TO SYNAPSE

REGULATION OF RECEPTOR LOCALIZATION TO SYNAPSE

REGULATION OF RESPIRATORY GASEOUS EXCHANGE BY NERVOUS SYSTEM PROCESS

REGULATION OF SHORT TERM NEURONAL SYNAPTIC PLASTICITY

REGULATION OF SMOOTHENED SIGNALING PATHWAY INVOLVED IN DORSAL VENTRAL NEURAL TUBE PATTERNING

REGULATION OF SPHINGOLIPID BIOSYNTHETIC PROCESS

REGULATION OF SYNAPSE ASSEMBLY

REGULATION OF SYNAPSE MATURATION

REGULATION OF SYNAPSE STRUCTURE OR ACTIVITY

REGULATION OF SYNAPTIC PLASTICITY

REGULATION OF SYNAPTIC TRANSMISSION GABAERGIC

REGULATION OF SYNAPTIC TRANSMISSION GLUTAMATERGIC

REGULATION OF SYNAPTIC VESICLE ENDOCYTOSIS

REGULATION OF SYNAPTIC VESICLE EXOCYTOSIS

REGULATION OF SYNAPTIC VESICLE RECYCLING

REGULATION OF TRANS SYNAPTIC SIGNALING

REGULATION OF TRANSMISSION OF NERVE IMPULSE

RESPONSE TO ACETYLCHOLINE

RESPONSE TO AMYLOID BETA

RESPONSE TO CORTICOSTEROID

RESPONSE TO CORTICOSTERONE

RESPONSE TO DOPAMINE

RESPONSE TO EPINEPHRINE

RESPONSE TO HISTAMINE

RESPONSE TO L GLUTAMATE

RESPONSE TO LEPTIN

RESPONSE TO MINERALOCORTICOID

RESPONSE TO NERVE GROWTH FACTOR

ROSTROCAUDAL NEURAL TUBE PATTERNING

SEMAPHORIN PLEXIN SIGNALING PATHWAY INVOLVED IN NEURON PROJECTION GUIDANCE

SEROTONIN METABOLIC PROCESS

SEROTONIN RECEPTOR SIGNALING PATHWAY

SEROTONIN TRANSPORT

SEROTONIN UPTAKE

SHORT TERM MEMORY

SKELETAL MUSCLE ACETYLCHOLINE GATED CHANNEL CLUSTERING

SLEEP

SMOOTHENED SIGNALING PATHWAY INVOLVED IN DORSAL VENTRAL NEURAL TUBE PATTERNING

SPHINGOLIPID BIOSYNTHETIC PROCESS

SPHINGOLIPID MEDIATED SIGNALING PATHWAY

SPHINGOLIPID METABOLIC PROCESS

SPHINGOMYELIN BIOSYNTHETIC PROCESS

SPHINGOMYELIN METABOLIC PROCESS

SPINAL CORD ASSOCIATION NEURON DIFFERENTIATION

SPINAL CORD MOTOR NEURON DIFFERENTIATION

SPONTANEOUS SYNAPTIC TRANSMISSION

SYMPATHETIC NERVOUS SYSTEM DEVELOPMENT

SYNAPSE ASSEMBLY

SYNAPSE MATURATION

SYNAPSE ORGANIZATION

SYNAPSE PRUNING

SYNAPTIC MEMBRANE ADHESION

SYNAPTIC SIGNALING

SYNAPTIC TRANSMISSION CHOLINERGIC

SYNAPTIC TRANSMISSION DOPAMINERGIC

SYNAPTIC TRANSMISSION GABAERGIC

SYNAPTIC TRANSMISSION GLUTAMATERGIC

SYNAPTIC VESICLE CLUSTERING

SYNAPTIC VESICLE CYTOSKELETAL TRANSPORT

SYNAPTIC VESICLE DOCKING

SYNAPTIC VESICLE EXOCYTOSIS

SYNAPTIC VESICLE LOCALIZATION

SYNAPTIC VESICLE MATURATION

SYNAPTIC VESICLE MEMBRANE ORGANIZATION

SYNAPTIC VESICLE PRIMING

SYNAPTIC VESICLE RECYCLING

SYNAPTIC VESICLE TRANSPORT

SYNAPTONEMAL COMPLEX ORGANIZATION

TELENCEPHALON GLIAL CELL MIGRATION

TRANS SYNAPTIC SIGNALING MODULATING SYNAPTIC TRANSMISSION

TRANSMISSION OF NERVE IMPULSE

TRIGEMINAL NERVE DEVELOPMENT

VENTRAL SPINAL CORD INTERNEURON DIFFERENTIATION

VESICLE MEDIATED TRANSPORT IN SYNAPSE

VESTIBULOCOCHLEAR NERVE DEVELOPMENT

WNT SIGNALING PATHWAY INVOLVED IN MIDBRAIN DOPAMINERGIC NEURON DIFFERENTIATION

**Supplementary Text Table 2.** Music-related genes from Navarro et al. [9] that are DE in TP2 vs. TP1 in ACD patients. All these genes are protein coding.

| **IDs** | **baseMean** | **log_2_FC** | **log_2_FC–SD** | **stat** | ***P*-value** | **FDR-value** | **Gene symbol** |
| --- | --- | --- | --- | --- | --- | --- | --- |
| ENSG00000002834 | 6219.8 | -0.2841 | 0.0558 | -5.0958 | 0.0000 | 0.0017 | *PABPC1* |
| ENSG00000004939 | 653.7 | 0.3496 | 0.0920 | 3.7979 | 0.0001 | 0.0164 | *ADRB2* |
| ENSG00000028116 | 334.4 | 0.2296 | 0.0624 | 3.6768 | 0.0002 | 0.0204 | *GZMK* |
| ENSG00000050130 | 670.6 | 0.3109 | 0.0866 | 3.5879 | 0.0003 | 0.0250 | *CASP3* |
| ENSG00000064012 | 787.1 | -0.3821 | 0.1074 | -3.5570 | 0.0004 | 0.0269 | *APOBR* |
| ENSG00000065978 | 738.7 | 0.2808 | 0.0830 | 3.3855 | 0.0007 | 0.0354 | *JKAMP* |
| ENSG00000068024 | 2627.1 | -0.1792 | 0.0532 | -3.3690 | 0.0008 | 0.0358 | *RNF213* |
| ENSG00000070756 | 762.2 | 0.1678 | 0.0527 | 3.1842 | 0.0015 | 0.0464 | *NPTN* |
| ENSG00000076944 | 365.0 | -0.2248 | 0.0708 | -3.1756 | 0.0015 | 0.0469 | *HDAC4* |
| ENSG00000088832 | 529.6 | 0.1910 | 0.0611 | 3.1251 | 0.0018 | 0.0496 | *TFG* |
| ENSG00000104695 | 125.0 | -0.2339 | 0.0764 | -3.0618 | 0.0022 | 0.0539 | *KCTD9* |
| ENSG00000104756 | 336.7 | 0.3604 | 0.1187 | 3.0365 | 0.0024 | 0.0554 | *CXCL8* |
| ENSG00000109332 | 664.1 | -0.3155 | 0.1058 | -2.9828 | 0.0029 | 0.0590 | *SLC6A8* |
| ENSG00000109756 | 1909.5 | 0.2181 | 0.0734 | 2.9715 | 0.0030 | 0.0606 | *CASP8* |
| ENSG00000111859 | 8816.9 | 0.1966 | 0.0697 | 2.8210 | 0.0048 | 0.0745 | *UBE2D3* |
| ENSG00000112893 | 2768.6 | 0.1311 | 0.0472 | 2.7792 | 0.0054 | 0.0798 | *DUSP6* |
| ENSG00000113088 | 3989.2 | -0.2338 | 0.0845 | -2.7666 | 0.0057 | 0.0816 | *YBX1* |
| ENSG00000114354 | 273.0 | 0.1846 | 0.0669 | 2.7617 | 0.0058 | 0.0819 | *ZNF83* |
| ENSG00000114861 | 580.4 | 0.1924 | 0.0701 | 2.7461 | 0.0060 | 0.0839 | *HIGD1A* |
| ENSG00000114978 | 2584.2 | 0.1378 | 0.0511 | 2.6984 | 0.0070 | 0.0889 | *CX3CR1* |
| ENSG00000115310 | 1146.9 | 0.1768 | 0.0658 | 2.6859 | 0.0072 | 0.0905 | *DSTN* |
| ENSG00000115758 | 103.1 | 0.2963 | 0.1117 | 2.6532 | 0.0080 | 0.0951 | *PKIA* |
| ENSG00000118260 | 1869.3 | 0.1857 | 0.0702 | 2.6475 | 0.0081 | 0.0951 | *ODC1* |
| ENSG00000125868 | 1777.4 | 0.2082 | 0.0806 | 2.5834 | 0.0098 | 0.1046 | *TLR1* |
| ENSG00000130821 | 104.5 | 0.3192 | 0.1252 | 2.5490 | 0.0108 | 0.1101 | *HDC* |
| ENSG00000135336 | 1342.0 | -0.1367 | 0.0547 | -2.4990 | 0.0125 | 0.1198 | *RTN4* |
| ENSG00000139318 | 1690.5 | 0.1506 | 0.0606 | 2.4843 | 0.0130 | 0.1230 | *SDHD* |
| ENSG00000140287 | 6115.3 | -0.1650 | 0.0678 | -2.4322 | 0.0150 | 0.1311 | *RPS9* |
| ENSG00000142197 | 944.2 | -0.1469 | 0.0610 | -2.4081 | 0.0160 | 0.1356 | *FOXP1* |
| ENSG00000156642 | 1866.8 | 0.2115 | 0.0881 | 2.4017 | 0.0163 | 0.1366 | *FOS* |
| ENSG00000164305 | 1188.8 | 0.1605 | 0.0670 | 2.3967 | 0.0165 | 0.1367 | *CREB1* |
| ENSG00000167766 | 287.1 | 0.1708 | 0.0713 | 2.3953 | 0.0166 | 0.1367 | *ORC3* |
| ENSG00000168329 | 242.4 | -0.1750 | 0.0731 | -2.3930 | 0.0167 | 0.1368 | *RAPGEF2* |
| ENSG00000169252 | 1099.2 | -0.1610 | 0.0693 | -2.3247 | 0.0201 | 0.1491 | *LASP1* |
| ENSG00000169429 | 166.5 | -0.1725 | 0.0748 | -2.3065 | 0.0211 | 0.1524 | *DOP1B* |
| ENSG00000170345 | 964.3 | 0.1352 | 0.0595 | 2.2714 | 0.0231 | 0.1598 | *NEDD9* |
| ENSG00000170889 | 164.5 | -0.2096 | 0.0932 | -2.2490 | 0.0245 | 0.1663 | *PPP2CB* |
| ENSG00000171033 | 589.1 | -0.1444 | 0.0644 | -2.2435 | 0.0249 | 0.1674 | *MAN2A1* |
| ENSG00000173821 | 2142.1 | 0.1478 | 0.0669 | 2.2075 | 0.0273 | 0.1765 | *MYADM* |
| ENSG00000174125 | 541.4 | -0.1757 | 0.0797 | -2.2058 | 0.0274 | 0.1769 | *FKBP1A* |
| ENSG00000179820 | 157.7 | 0.1906 | 0.0877 | 2.1726 | 0.0298 | 0.1857 | *VRK2* |
| ENSG00000181061 | 3247.8 | 0.1147 | 0.0536 | 2.1397 | 0.0324 | 0.1929 | *MOB1A* |
| ENSG00000183813 | 1246.7 | -0.1112 | 0.0542 | -2.0535 | 0.0400 | 0.2148 | *STXBP2* |
| ENSG00000184678 | 454.6 | 0.1894 | 0.0934 | 2.0282 | 0.0425 | 0.2208 | *H2BC21* |
| ENSG00000184730 | 2225.6 | -0.1612 | 0.0797 | -2.0241 | 0.0430 | 0.2214 | *SLC4A1* |
| ENSG00000204370 | 146.3 | -0.2153 | 0.1073 | -2.0069 | 0.0448 | 0.2253 | *PLIN5* |
| ENSG00000211456 | 553.9 | 0.1334 | 0.0680 | 1.9620 | 0.0498 | 0.2394 | *SACM1L* |
| ENSG00000214456 | 86.4 | 0.2526 | 0.1285 | 1.9658 | 0.0493 | NA | *CCR4* |
